# Reinforcement learning-driven unified generative framework for multi-objective RNA codon design

**DOI:** 10.64898/2026.06.12.732012

**Authors:** Shenggeng Lin, Hong Tan, Keyao Wang, Ruixuan Wang, Hongxia Wang, Tong Zhu, Yi Xiong

**Author notes:** Corresponding author(s): Yi Xiong or Tong Zhu.

## Abstract

Current RNA codon design methods are limited by inefficient long-sequence processing and poor generalizability, often relying on a decoupled ‘generate-or-optimize’ paradigm. We introduce RNARL, a reinforcement learning-driven framework that unifies sequence generation with multi-objective optimization. RNARL directly learns to generate high-performance sequences, effectively optimizing sequences over 3,900 nucleotides and demonstrating superior performance and universality across six species and five RNA types. RNARL thus establishes an effective and generalizable framework for RNA codon design. Finally, a user-friendly web platform is freely available to facilitate its application for RNA therapeutic design.

## Background

RNA vaccines deliver RNA that encodes antigens, enabling host cells to autonomously synthesize these antigens and activate immune responses [1–4]. However, due to the degeneracy of the genetic code, a single protein sequence can be encoded by multiple distinct RNA sequences [5, 6]. This codon degeneracy poses a critical challenge: although encoding identical proteins, synonymous RNA variants can exhibit significant differences in translational efficiency, structural stability and expression levels [7]. These differences primarily stem from species-specific codon usage bias and RNA structural features [8, 9]. Therefore, RNA codon design constitutes a critical strategy for achieving optimal vaccine efficacy [10–14]. This process enhances sequence compatibility with host translational machinery while stabilizing RNA secondary structures, thereby amplifying antigen expression to potentiate both prophylactic and therapeutic applications [15–17].

Despite the demonstrated importance of codon design [18], current implementation strategies encounter multifaceted challenges. Traditional strategies, such as ‘host-preferred codon substitution’ and ‘natural codon distribution matching’, are conceptually straightforward but primarily optimize for translational efficiency, often at the expense of RNA structural stability [19, 20]. These methods operate by replacing codons based on host frequency statistics. For example, the ‘host-preferred substitution’ strategy uses only the single most frequent codon for each amino acid. This emphasis on codon frequency inherently neglects the sequence-specific nucleotide context that is essential for optimal folding, often resulting in suboptimal trade-offs between translational efficiency and RNA structural stability. Moreover, such frequency-based codon designs can lead to the depletion of specific tRNA pools, further limiting their practical utility [5]. More recent approaches, such as LinearDesign [15], aim to achieve a balance between translational efficiency and structural stability, demonstrating promising solutions. However, their reliance on heuristic search strategies limits their efficiency in achieving high-throughput optimization of long RNA sequences. Deep learning-based methods [21–26], including those employing LSTM architectures, struggle to capture long-range dependencies in RNA sequences and are unsuitable for handling exceptionally long sequences [27]. Furthermore, pretrained models, often trained on natural sequence distributions, tend to generate sequences whose properties closely resemble their natural counterparts, thereby hindering their capacity to produce RNA sequences with substantially superior performance [5, 28–33].

Based on the aforementioned facts, current RNA codon design algorithms encounter three unresolved challenges: First, efficiently processing and optimizing long RNA sequences (>3,000 nucleotides) remains a critical bottleneck [15]. Second, existing methods demonstrate limited generalizability across species and RNA types [27]. Third, while generative models enable high throughput design, they rarely produce RNAs with properties superior to natural RNA sequences. In contrast, optimization algorithms can generate sequences that surpass natural ones, but are often too computationally intensive for large-scale design. The conventional ‘generate-or-optimize’ paradigm imposes inherent efficiency and performance limitations, necessitating deep integration of sequence generation with optimization objectives to directly generate high-performance sequences.

To address these three challenges, we propose RNA sequence design via reinforcement learning (named RNARL), a unified framework that integrates a mixture-of-experts (MoE) Transformer-based generative model [34] with group relative policy optimization (GRPO) [35] for RNA codon design. The core idea is to leverage the high-throughput sequence generation capabilities of generative models, while employing reinforcement learning to iteratively refine the generative policy in accordance with predefined multi-objective design criteria. In RNARL, the generative model is directly optimized based on a reward function that evaluates desired properties (e.g., high translational efficiency, enhanced structural stability), thereby refining its policy to generate RNA sequences that satisfy these objectives. This approach tightly couples sequence generation with optimization, distinguishing it from the conventional ‘generate-or-optimize’ paradigm and enabling the direct design of high-performance RNA sequences.

Extensive computational experiments validate the framework’s efficacy and universality. Notably, by training on datasets containing RNA sequences up to ∼3,900 nucleotides (nts) in length and incorporating length-aware distribution sampling (LADS) techniques, our framework overcomes existing limitations in processing long sequences, enabling efficient generation and optimization. Furthermore, evaluations across six major biological categories demonstrate its exceptional adaptability to diverse host environments and heterologous expression scenarios, while assessments on five RNA types confirm its broad applicability to diverse RNA molecular designs. Successful optimization of two publicly available RNA vaccine patent sequences by RNARL underscores the framework’s practical potential.

## Results

### Overview of the RNARL Framework

RNARL integrates the GRPO [35] reinforcement learning algorithm with generative models to optimize RNA sequences that encode target proteins. The framework comprises an actor model and a reward model (Fig. 1a). The actor model serves as a generator, taking protein sequences as input and producing candidate RNA sequences. The reward model acts as an evaluator, taking an RNA sequence as input and assigning it a quantitative score. The two models are trained jointly via reinforcement learning: the reward model defines a multi-objective reward signal, and the parameters of the actor model are iteratively updated to refine its generative policy, thereby favoring the generation of RNA sequences that maximize this reward.

**Fig. 1.**
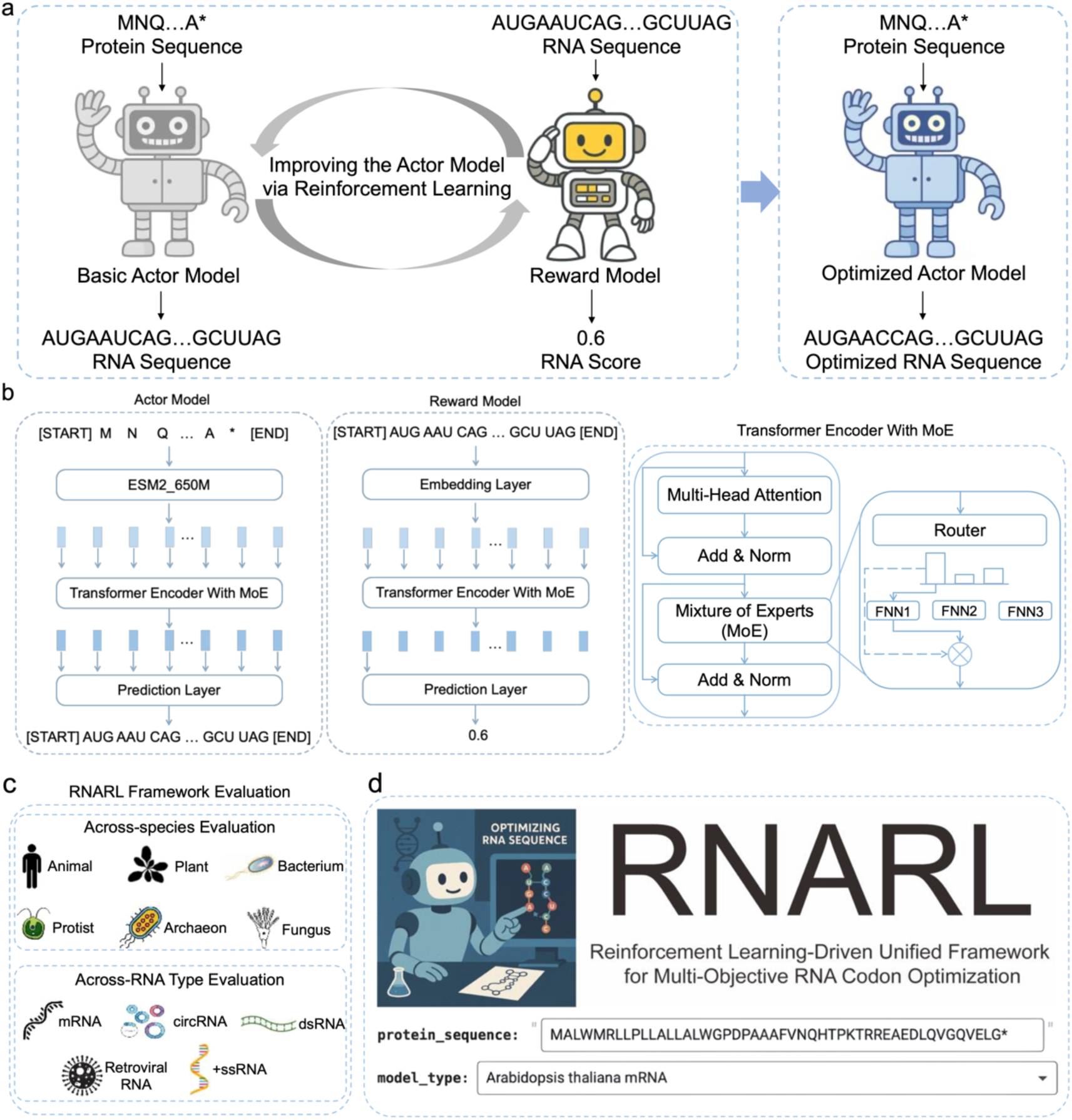
Overview of the RNARL framework. **a**, The reinforcement learning optimization process, where an actor model (generator) is iteratively refined based on reward signals from a reward model (evaluator). **b**, The core model architecture incorporating Transformer and MoE layers. **c**, Validation of the RNARL framework’s generalizability across six species and five RNA types. **d**, The user-friendly RNARL online design platform.

Both the actor and reward models are based on a Transformer encoder architecture enhanced with MoE layers (Fig. 1b). The framework’s performance was rigorously evaluated across six diverse biological categories: *Homo sapiens* (animals), *Arabidopsis thaliana* (plants), *Penicillium chrysogenum* (fungi), *Chlamydomonas reinhardtii* (protists), *Escherichia coli* (bacteria), and *Thermococcus kodakarensis* KOD1 (archaea). Its generalizability was further tested on five distinct RNA types, including mRNA, circular RNA (circRNA), positive-sense single-stranded RNA (+ssRNA), double-stranded RNA (dsRNA), and retroviral RNA (Fig. 1c). Furthermore, a user-friendly web platform was developed to facilitate RNA therapeutic design (Fig. 1d).

### Dataset Benchmarking and Ablation Analysis

To comprehensively evaluate the capabilities of the RNARL framework, we constructed six datasets spanning distinct biological categories. These datasets cover protein lengths ranging from 10 to 1,305 amino acids (corresponding to RNA lengths of 30–3,915 nt), encompassing most potential antigen length ranges. Each dataset was rigorously partitioned into training, validation, and test sets (Fig. 2a, see Materials and Methods). We performed detailed analyses across the datasets and present the *Homo sapiens* dataset as an illustrative example of our findings.

**Fig. 2.**
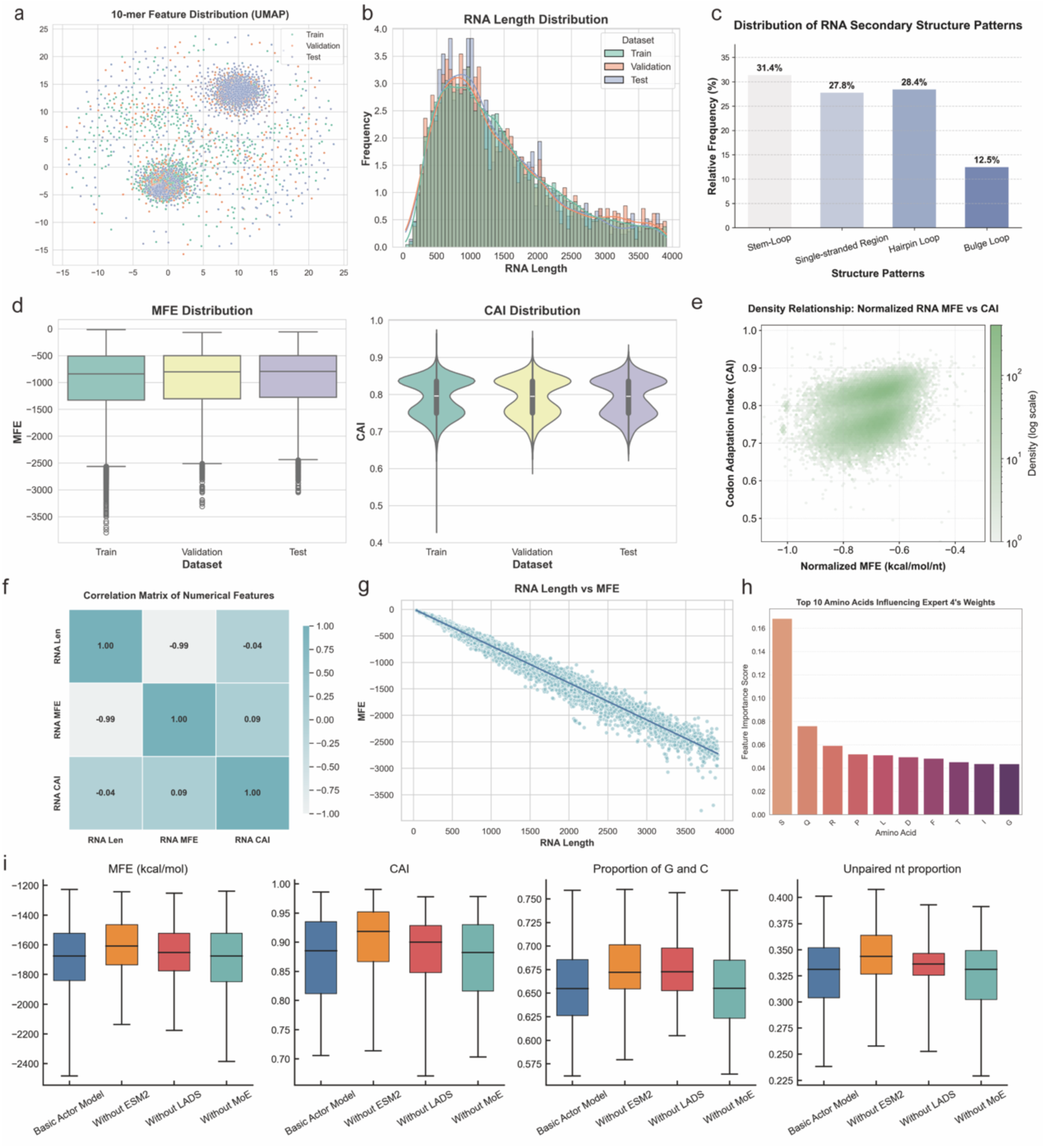
Characterization of the *Homo sapiens* dataset and ablation study of the RNARL framework. **a**, Visualization of distribution of protein sequences using 10-mer frequency vectors across the training, validation, and test sets. **b**, The distribution of RNA lengths (in nucleotides) for sequences on the training, validation, and test datasets. **c,** The relative frequency of different RNA secondary structure patterns (Stem-Loop, Single-stranded region, Hairpin Loop, Bulge Loop) within the dataset. **d,** Distributions of MFE and CAI for RNA sequences across the training, validation, and test sets. **e,** The joint distribution of normalized MFE and CAI values; color intensity indicates density. **f,** Pearson correlation coefficients between RNA length (RNA Len), RNA MFE, and RNA CAI values. **g,** The correlation between RNA length and MFE, with a fitted polynomial regression line. **h,** The top 10 amino acid types influencing the activation status (weights) of Expert 4 in the MoE gating network, based on feature importance scores. **i,** Ablation experiments on the RNARL framework, comparing the performance of the basic actor model with variants lacking key components (without ESM2, without LADS, without MoE).

A pronounced ‘long-tail distribution’ was observed in RNA sequence lengths: when binned into 100-nt intervals, sequence counts varied by up to an order of magnitude across intervals (Fig. 2b). This imbalance poses challenges for existing models in effectively optimizing long sequences. To mitigate this issue, we employed LADS techniques during the model training. RNA structural features play a crucial role in RNA structural stability. We quantified the frequency of major RNA secondary structures in the dataset (Fig. 2c). Stem-loop and hairpin loop structures were dominated, accounting for 31.4% and 28.4% of the total ones, respectively, consistent with their roles in maintaining RNA stability and regulating translation processes.

Four core metrics were used to evaluate the generated RNA sequences: Minimum Free Energy (MFE; lower values correlate with higher structural stability), Codon Adaptation Index [36] (CAI; higher values correlate with enhanced expression potential), proportion of G and C [37] (a measure of structural stability; higher proportion of G and C may favor stable secondary structures), and unpaired nt proportion (a measure of flexibility/degradation risk; lower values reduce susceptibility to ribonuclease degradation). We further analyzed two key optimization metrics, that are MFE and CAI, for RNA sequences on the dataset. Overall, MFE and CAI distributions were similar across subsets (Fig. 2d). Joint analysis of CAI and normalized MFE revealed a weak positive correlation (r = 0.333, p < 0.01; Fig. 2e), consistent with a trade-off between RNA structural stability (lower MFE) and translational efficiency (higher CAI), which poses a key challenge for RNA design algorithms. Additionally, a negative correlation between RNA length and MFE supports the use of normalized MFE (MFE divided by sequence length) as a length-agnostic stability metric (Fig. 2f, g).

On the *Homo sapiens* dataset, hyperparameter tuning demonstrated the robustness of the RNARL model to parameter variations (Supplementary Table 1). Ablation experiments showed that both the ESM-2 [38] encoder and the LADS strategy are essential for multi-objective optimization in RNARL (Fig. 2i). Removal of either component results in over-optimization for translation efficiency at the expense of structural stability, leading to an improved CAI but also an increased MFE and unpaired nt proportion. Analysis of the MoE [39] gating network revealed its ability to activate specific experts based on input sequence features, enhancing the model’s adaptability to diverse protein characteristics (Supplementary Note 1, Supplementary Fig. 1). For instance, Expert 4 specializes in processing serine-rich sequences or serine-associated patterns (Fig. 2h).

### Cross-Species RNA Codon Design Capacity of the RNARL Framework

To validate the universality of the RNARL framework across species, we trained six species-specific models based on the RNARL framework using the previously described datasets. The performance comparison between a unified model and six species-specific models is presented in Supplementary Note 9 and Supplementary Fig. 10. RNARL-generated RNA sequences were benchmarked against five methods (GEMORNA [32], LinearDesign [15], CodonTransformer [5], CAI-optimized, and Random) and natural sequences from all six species (Fig. 3, Supplementary Fig. 2). Specifically, GEMORNA and CodonTransformer represent advanced approaches based on multi-species pre-trained large language models; LinearDesign represents a state-of-the-art heuristic search method; CAI-optimized represents a traditional codon optimization method; and Random servers as the baseline.

**Fig. 3.**
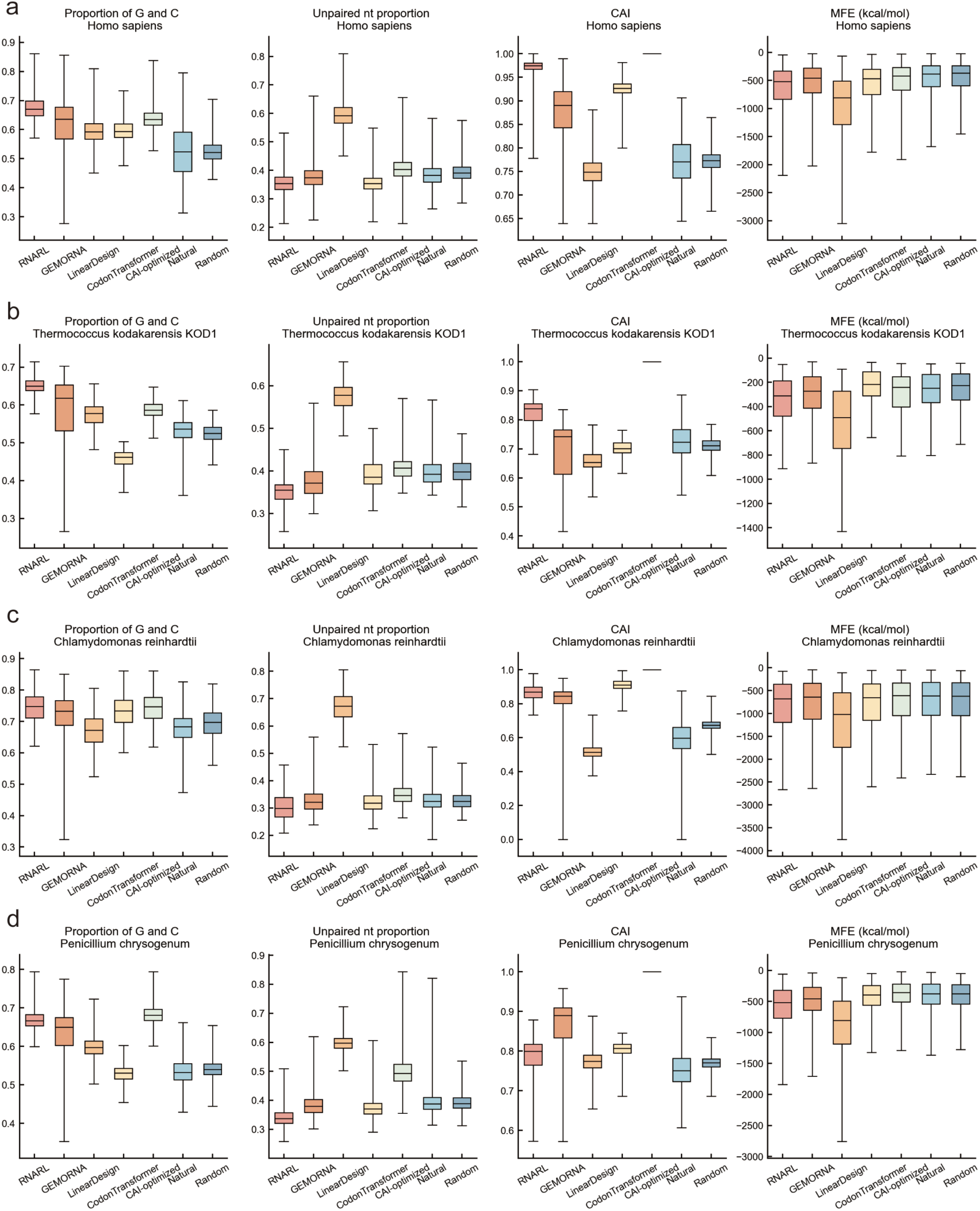
Cross-species evaluation of codon design methods. The codon design methods compared include GEMORNA, RNARL, LinearDesign, Codon Transformer, CAI-optimized, Random sequences, and Natural (representing the wild-type RNA sequence). **a,** Results for *Homo sapiens*. **b,** Results for *Thermococcus kodakarensis KOD1*. **c,** Results for *Chlamydomonas reinhardtii*. **d,** Results for *Penicillium chrysogenum*.

RNARL outperformed other methods and natural sequences across multiple metrics, including proportion of G and C, unpaired nt proportion, and CAI metrics. For example, on *Homo sapiens*, RNARL-generated RNA sequences exhibited the proportion of G and C of 0.676 ± 0.040, compared to 0.617 ± 0.090 for GEMORNA, 0.596 ± 0.045 for LinearDesign and 0.598 ± 0.036 for CodonTransformer. RNARL also achieved a CAI of 0.967 ± 0.027, outperforming GEMORNA’s CAI of 0.877 ± 0.057, LinearDesign’s CAI of 0.749 ± 0.030 and CodonTransformer’s CAI of 0.926 ± 0.015. Although the CAI-optimized method attained the maximum CAI scores (1.0) through optimal codon selection, it demonstrated inferior performance across other metrics. In contrast, RNARL maintained high CAI values while achieving a more balanced performance across all evaluated metrics. LinearDesign showed lowest (optimal) MFE values, while RNARL exhibited marginally higher (less optimal) MFE values than LinearDesign but still outperformed all other methods. The superior MFE of LinearDesign-generated sequences is expected, as LinearDesign is a heuristic search algorithm that can, in principle, find the lowest MFE sequence when its lambda hyperparameter is set to 0 and the search width is sufficiently large. In addition, a comparison of sequence similarity revealed that RNA sequences generated by both RNARL and CodonTransformer exhibit >70% similarity to their corresponding natural CDS (Supplementary Note 7, Supplementary Fig. 8). Overall, RNARL’s reinforcement learning-driven multi-objective design achieved well-balanced performance across all metrics, demonstrating its effectiveness in generating high-quality RNA sequences.

To quantify the influence of protein properties on RNARL performance, we analyzed the relationship between the optimization gains, defined as normalized MFE Gain and CAI Gain, and the intrinsic physicochemical properties of the corresponding proteins. A strong positive correlation between normalized MFE Gain and CAI Gain (Spearman’s r = 0.64, p < 0.01, Supplementary Fig. 11a, b) indicates that RNARL can simultaneously enhance RNA stability and translational efficiency, underscoring its capability for multi-objective optimization. Furthermore, ‘easy-to-optimize’ proteins are more hydrophobic and exhibit lower instability indices, whereas ‘hard-to-optimize” proteins are less hydrophobic and more unstable (Supplementary Fig. 11f). A detailed description of the methodology and extended analyses are provided in Supplementary Note 10.

T-SNE visualization of the protein sequence embeddings derived from the actor model revealed species-specific clustering, with analogous patterns observed in RNARL-generated RNA sequences (Fig. 4a, b). These results confirm that RNARL effectively learns host-specific codon preferences and preserves phylogenetically conserved sequence features. Codon pairs have been demonstrated to significantly influence protein expression levels by modulating translational kinetics [40, 41]. Quantitative analysis of undesirable codon pair frequencies demonstrated that RNARL and LinearDesign generated sequences with fewer inhibitory codon pairs compared to other methods, supporting their enhanced translation efficiency (Fig. 4c) [42]. While thermophilic organisms typically exhibit high proportion of G and C in genomic DNA to enhance thermostability, excessive GC pairs in RNA may form overly stable secondary structures that hinder ribosomal translocation [43–45]. RNARL selectively suppressed GC-rich codon pairs of *Thermococcus kodakarensis KOD1*. The proportion of GC-rich codon pairs in RNARL-generated RNA sequences was comparable to those in LinearDesign-generated sequences and 36.7% lower than that in natural sequences, potentially mitigating the formation of RNA secondary structures. This adaptive codon optimization strategy mirrors natural evolutionary pressures in hyperthermophiles, where reduced GC content in coding sequences minimizes thermodynamically stable RNA structures that could impede ribosome progression during translation at elevated temperatures [46, 47]. To validate heterologous expression compatibility, the RNARL model trained on *Homo sapiens* data was utilized to optimize RNA sequences encoding *E. coli* proteins. Remarkably, the CAI (calculated based on human codon usage preferences) of *E. coli* RNA sequences optimized by the *Homo sapiens*-trained RNARL model was approximately 0.182 higher than that of sequences optimized by the *E. coli*-trained RNARL model, while maintaining comparable structural stability metrics. This demonstrates RNARL’s robust alignment with target host codon preferences during cross-species optimization (Fig. 4d). These results highlight the versatility of the RNARL framework in RNA codon optimization across diverse species.

**Fig. 4.**
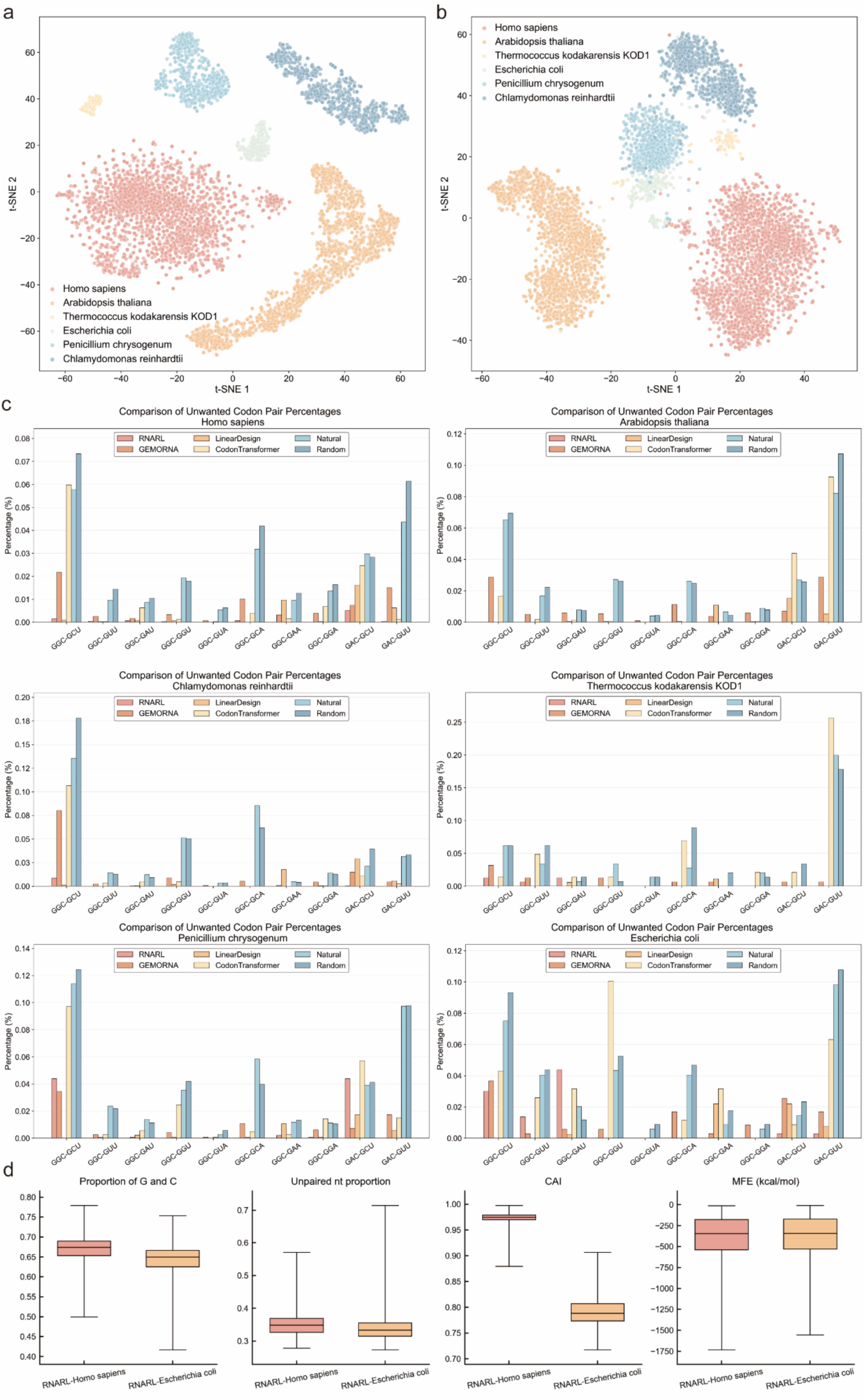
The capability of RNARL for cross-species RNA sequence representation and design. **a.** t-SNE visualization of protein sequence embeddings extracted using the actor model. Each point represents a protein sequence, colored by its species origin, demonstrating clear species-specific clustering. **b.** t-SNE visualization of RNARL-generated RNA sequences using 4-mer frequency vectors. Each point represents an RNA sequence, colored by its species origin, demonstrating clear species-specific clustering. **c.** Comparison of undesirable codon pair frequencies in RNA sequences optimized by RNARL versus other methods across six species. **d.** Evaluation of RNA sequences optimized by the RNARL-*Homo sapiens* model for *Escherichia coli* RNAs (cross-species optimization) and the RNARL-*Escherichia coli* model for *Escherichia coli* RNAs (intra-species optimization).

### Cross-Type RNA Codon Design Capacity of the RNARL Framework

While mRNA remains the predominant focus for vaccine development, other RNA types exhibit substantial therapeutic potential in gene therapy, immune modulation, and innovative vaccine design, owing to their distinct biological characteristics (e.g., circular RNA) [48–50]. To systematically validate the generalizability of the RNARL framework across RNA types, we further evaluated its performance on four additional RNA types: circRNA, +ssRNA, dsRNA, and retroviral RNA. These RNA types differ significantly from linear mRNA in both structure and function, thereby posing unique challenges and requirements for optimization strategies. For instance, circRNA exhibits inherent resistance to nuclease degradation due to its closed-loop structure and absence of free termini, making it an ideal vector for sustained expression. +ssRNA and retroviral RNA, as viral genomes or intermediates, rely critically on their structural and sequence features for replication, gene expression, and packaging.

As shown in Fig. 5a, the results indicate that RNARL-generated sequences consistently outperformed other methods in circRNA optimization with respect to the proportion of G and C, unpaired nt proportion, and CAI compared to other methods. For instance, RNARL-generated sequences exhibited the proportion of G and C of 0.661 ± 0.047, compared to 0.588 ± 0.111 (GEMORNA), 0.579 ± 0.054 (LinearDesign) and 0.584 ± 0.040 (CodonTransformer), respectively. The optimized proportion of G and C and a favorable unpaired nt proportion are vital for the overall stability and functional integrity of circRNA, as well as for enhanced resistance to enzymatic degradation [51–53]. Furthermore, across the three viral RNA types (+ssRNA, dsRNA, and retroviral RNA), RNARL-generated sequences consistently demonstrated superior performance in both proportion of G and C and CAI compared to other methods (Fig. 5b, c, Supplementary Fig. 3). For +ssRNA, the CAI of RNARL-generated sequences was approximately 0.104 higher than that of GEMORNA-generated sequences, 0.239 higher than that of LinearDesign-generated sequences and 0.043 higher than that of CodonTransformer-generated sequences. A higher CAI is essential for efficient viral replication and robust protein expression in viral vectors [54–56]. For +ssRNA, elevated GC content and reduced UA/UU dinucleotide content significantly mitigate the likelihood of degradation by RNase L. For instance, a comparative analysis of the wild-type RNA sequence encoding the AAT98583.1 protein of Human coronavirus HKU1 and its RNARL-generated counterpart revealed that the wild-type sequence contained a substantially higher abundance of UA and UU dinucleotides, which are prone to RNase L cleavage, rendering it more susceptible to RNase L-mediated degradation (Fig. 5d).

**Fig. 5.**
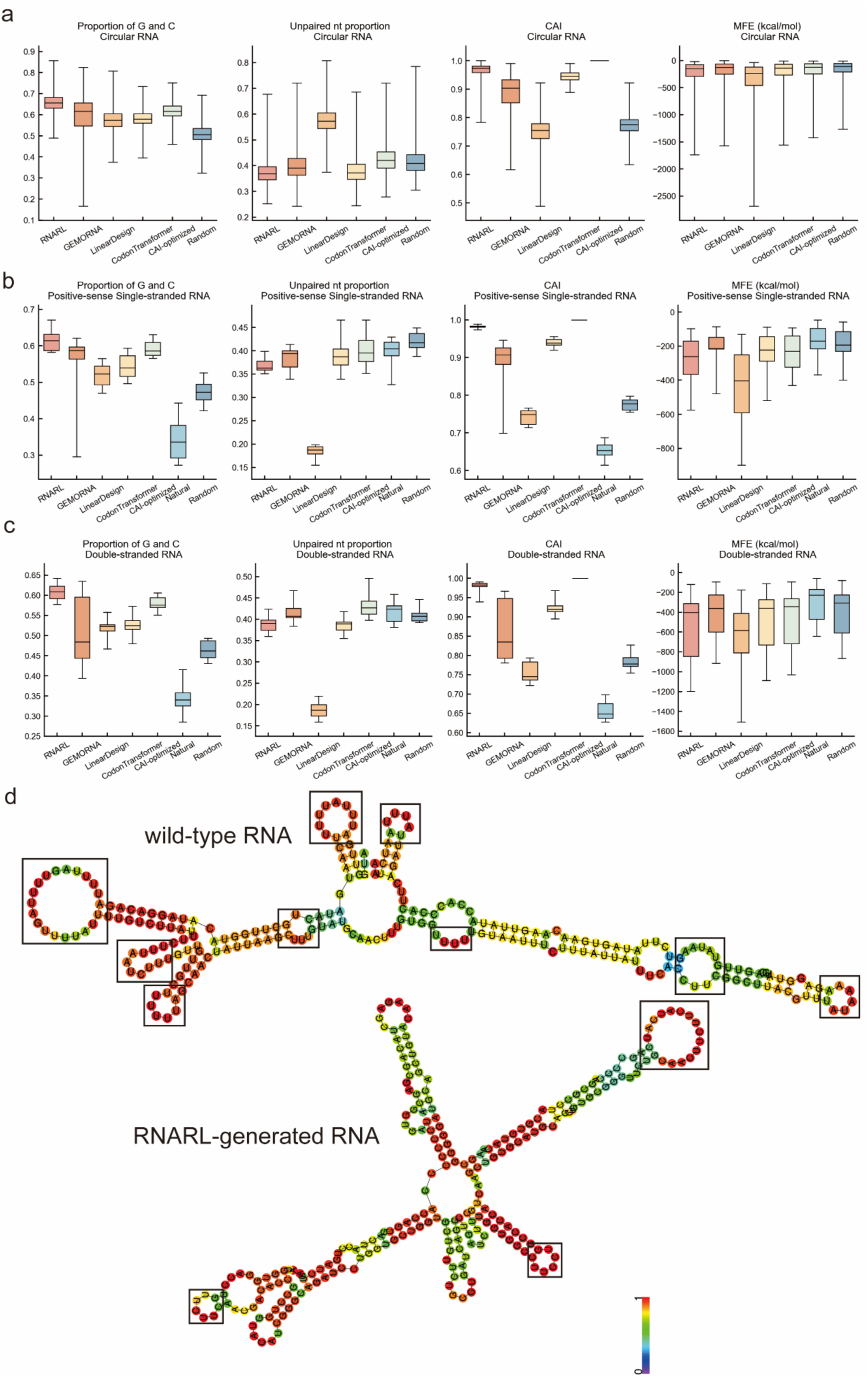
Performance evaluation of codon design methods across diverse RNA types. **a,**Results for circRNA. **b**, Results for positive-sense single-stranded RNA. **c**, Results for double-stranded RNA. **d,** Secondary structure comparison and RNase L susceptibility. Predicted secondary structure of the wild-type RNA sequence encoding human coronavirus HKU1 protein AAT98583.1, compared to the corresponding sequence generated by RNARL. Regions indicated by black boxes are susceptible to RNase L-mediated degradation. The color of each nucleotide reflects its base-pairing probability, where red indicates a high probability of being paired and blue indicates a low probability.

In summary, these results underscore the RNARL framework’s robust and generalizable optimization capacity across diverse RNA types. By consistently improving key metrics such as proportion of G and C, CAI, and structural stability, RNARL effectively addresses the distinct biological requirements of each RNA type, resulting in enhanced functional properties such as translational efficiency, structural stability, and nuclease resistance.

### Application of the RNARL Framework in RNA Vaccine Design

To demonstrate the utility of our RNARL framework in RNA vaccine design, we redesigned antigen-encoding RNA sequences derived from two clinically relevant RNA vaccine patents: (1) an mRNA vaccine targeting pathogenic gene regions of human papillomavirus (HPV) types 16 and 18, connected by a flexible peptide linker (Patent No.: CN116019904A); and (2) an mRNA vaccine encoding four critical glycoproteins (gP220, gP42, gL, gH) of the Epstein-Barr virus (EBV) (Patent No.: CN117529335A) (Fig. 6a).

**Fig. 6.**
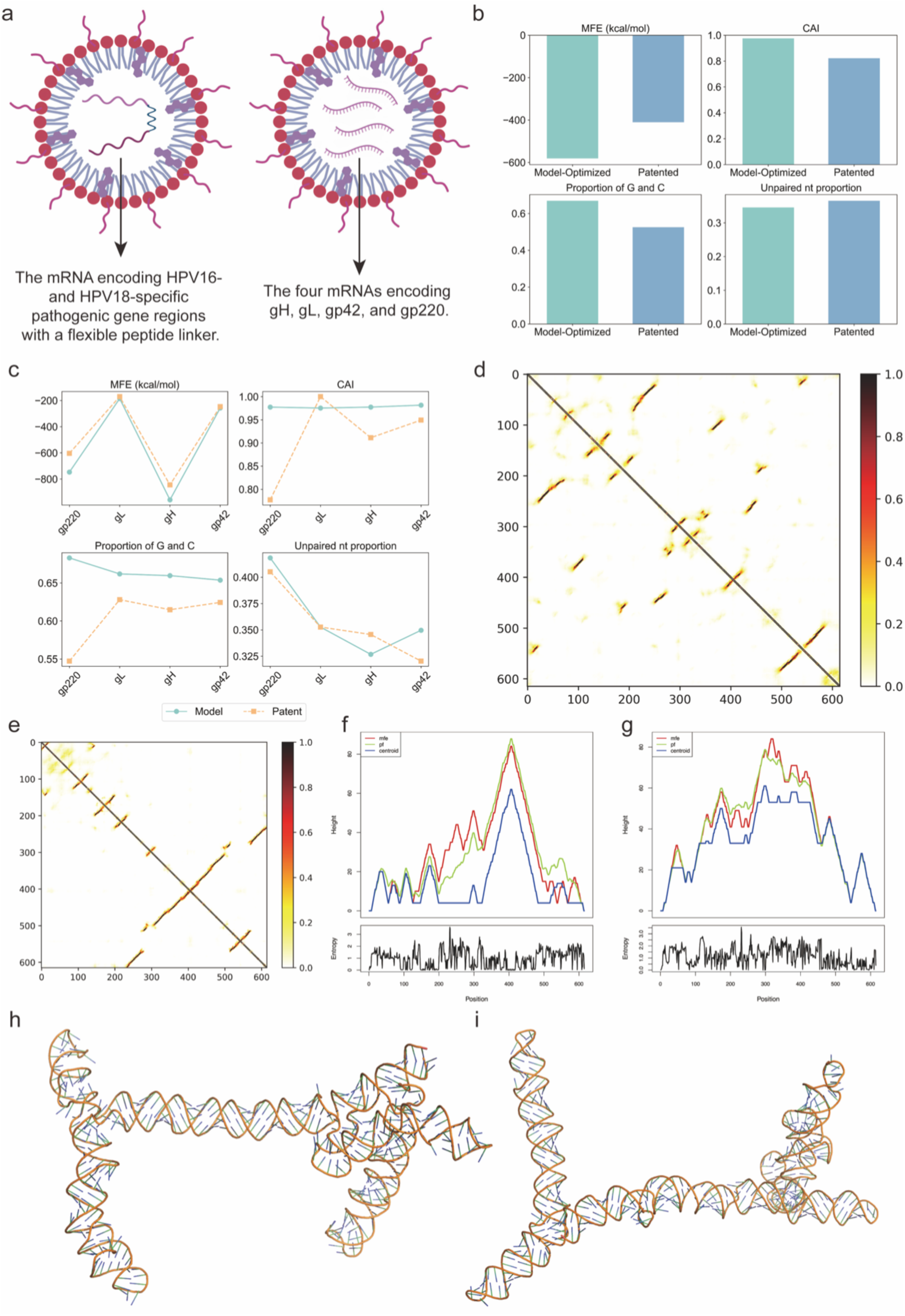
Structural and biophysical evaluation of patented and RNARL-optimized mRNA vaccine designs. **a,** Schematic diagrams illustrating the two types of mRNA vaccines. The vaccine described in patent CN116019904A contains a single mRNA encoding concatenated pathogenic gene regions of HPV types 16 and 18. The vaccine from patent CN117529335A contains four distinct mRNA sequences, each encoding one of four critical glycoproteins (gP220, gP42, gL, gH). **b, c,** Comparison of key RNA properties between RNARL-optimized and patented RNA sequences for the HPV vaccine (b) and the EBV vaccine (c). **d, e,** Structural contact maps of the RNA sequence encoding gp42 for the RNARL-optimized design (d) and the patented design (e). **f, g,** Comprehensive secondary structure analysis of the gp42-encoding RNA sequences for the patented (f) and RNARL-optimized design (g). Each panel contains two plots: (Top) Mountain plot representations of the secondary structure, where ‘mfe’ (red) is the MFE structure, ‘centroid’ (blue) is the most representative structure of the folding ensemble, and ‘pf’ (green) represents the average structure derived from the partition function calculation. (Bottom) Positional entropy, where higher values indicate greater structural uncertainty. **h, i,** Tertiary structures of the RNA sequence encoding gL for the patented design (h) and the RNARL-optimized design (i).

Using the RNARL framework, we generated optimized mRNA sequences for the antigen proteins derived from these patents. We then computationally compared these optimized sequences against their patent-provided counterparts. For the HPV vaccine, RNARL-optimized sequences consistently outperformed the patent sequences across all four key computational metrics (Fig. 6b). The MFE of the RNARL-generated sequence was -580.8 kcal/mol, compared to -410.4 kcal/mol for the patent RNA sequence. The CAI of the RNARL-generated sequence was 0.669, versus 0.526 for the patent RNA sequence. Similarly, for the EBV vaccine, the RNARL-generated sequences for gL and gH demonstrated superior or comparable performance in MFE, CAI, and proportion of G and C when compared to the patent sequences (Fig. 6c).

Beyond computational metrics, we conducted further structural analyses of the RNARL-optimized sequences for gL and gP42 antigens from the EBV vaccine patent (limited by the length constraints of current structural analysis tools). For gP42-encoding mRNAs, structural contact maps showed a higher number of contact points in RNARL-optimized sequences, suggesting the formation of more compact and stable tertiary structures (Fig. 6d-e, Supplementary Note 2, Supplementary Fig. 4). A comparative analysis of tertiary structures of gL-encoding mRNAs revealed that the RNARL-optimized sequence (Fig. 6i) folds into a more compact and tightly packed conformation compared to the patent-provided sequence (Fig. 6h), particularly at the central junction of the optimized structure. In complex cellular environments, such enhanced structural stability enables RNA to better maintain its functional conformation, reduce nonspecific interactions, and potentially improve overall stability. The secondary structure and detailed visualization of the energy landscape further confirm these findings. While specific regions of the patent gP42-encoding mRNA sequence (e.g., 300–450 nt positions) exhibited significant structural energy fluctuations (Fig. 6f), RNARL-optimized sequences maintained significantly higher structural stability in these critical regions (Fig. 6g). These intrinsically unstable regions in the patent sequences could serve as vulnerable targets for nuclease attacks in vivo, leading to reduced mRNA expression levels and duration.

In summary, the comprehensive evaluations demonstrate that the RNARL framework effectively generates RNA codon sequences with superior biophysical properties and enhanced structural stability across multiple dimensions. These strongly strongly validate the practical value and broad application potential of the RNARL framework in accelerating the rational design of high-quality RNA vaccine sequences.

## Discussion

The RNARL framework presented in this study addresses a fundamental limitation in RNA codon design: the decoupled ‘generate-or-optimize’ paradigm. Existing methods often excel at either high-throughput sequence generation or computationally intensive optimization, but struggle to achieve both goals simultaneously. RNARL bridges this gap by deeply unifying these two stages through reinforcement learning, enabling high-throughput generation of high-performance RNA sequences. Furthermore, it is worth emphasizing that RNARL is designed as a universal computational framework rather than a pretrained large model. The framework demonstrates high flexibility and modularity, and researchers can readily replace or adjust components such as the actor model, reward model, or reinforcement learning algorithms according to specific experimental or design requirements.

RNARL addresses at least three critical challenges in RNA codon design. First, by training on long RNA sequence datasets and incorporating LADS, RNARL effectively handles RNA sequences up to ∼3,900 nt in length. Second, computational validation across six distinct species and five RNA types demonstrates RNARL’s broad applicability, which is crucial for developing RNA therapies tailored to diverse host environments. Third, unlike single-objective methods, RNARL leverages reinforcement learning to more effectively jointly optimize multiple objectives, such as translation efficiency and structural stability, directly generating RNA sequences with enhanced overall performance.

Despite its strengths, the current RNARL framework has certain limitations. Its primary focus remains on coding sequence (CDS) optimization, while non-coding regions of RNA molecules (e.g., 5’/3’ UTRs of mRNA, IRES elements) play equally critical roles in RNA stability, translation efficiency, and subcellular localization. Given the high diversity of sequence and structural features in non-coding regions across RNA types, their unified incorporation into the design model presents complex challenges. Future work could explore the development of modular optimization components that design specific optimization strategies tailored to specific non-coding regions, which could then be integrated into the RNARL framework to enable global optimization of full-length RNA sequences.

### Conclusions

This study introduces and validates RNARL, a reinforcement-learning-driven framework for unified RNA sequence generation and multi-objective optimization. The results demonstrate that RNARL offers a practical and generalizable solution for designing RNA codon sequences with improved biophysical properties, such as enhanced translational efficiency and structural stability. The framework effectively handles long sequences across diverse species and RNA types.

By efficiently exploring the vast RNA sequence space, the RNARL framework holds the potential to accelerate the rational design of high-quality RNA vaccines and nucleic acid drugs. Its demonstrated ability to improve upon patented vaccine sequences highlights its translational relevance. Finally, the deployment of RNARL via a user-friendly web platform provides the community with an accessible tool to support future advances in the development of RNA-based medicine.

## Methods

### Dataset Construction and Preprocessing

To validate the generality and effectiveness of the RNARL framework across species and RNA types, we constructed datasets encompassing multi-species and multi-type RNA sequences.

#### Across-species mRNA Dataset

We downloaded and extracted the protein sequences and corresponding CDSs of six representative organisms from the National Center for Biotechnology Information (NCBI) database [57]: *Chlamydomonas reinhardtii* (protist), *Thermococcus kodakarensis* KOD1 (archaeon), *Homo sapiens* (animal), *Arabidopsis thaliana* (plant), *Escherichia coli* (bacterium), and *Penicillium chrysogenum* (fungus). To build a high-quality, non-redundant dataset, we clustered protein sequences using the Linclust algorithm [58]. Parameters were set to a minimum sequence identity threshold of 90% and a coverage threshold of 80%, ensuring that 80% of the length of one sequence aligned with another at 90% identity. One representative protein sequence and its corresponding RNA sequence were selected from each cluster to minimize bias from sequence homology. Sequences were filtered to retain proteins with lengths between 10 and 1,305 amino acids (corresponding to RNA lengths of 30–3,915 nucleotides), covering most potential antigen length ranges. Each species dataset was randomly partitioned into training set (90%), validation set (5%), and test set (5%). Detailed sequence counts for each species are provided in Supplementary Table 2. Codon usage bias information for each species was referenced from the Kazusa Codon Usage Database (https://www.kazusa.or.jp/codon/). The impact of sequence clustering identity thresholds on model performance is presented in Supplementary Note 8, Supplementary Fig. 9 and Supplementary Table 3.

#### Circular RNA Dataset

Human circRNAs and their corresponding protein sequences were downloaded from the circRNADb database [59]. Data processing followed a similar pipeline to the mRNA dataset, including clustering-based redundancy reduction, length filtering and dataset partitioning.

#### Viral RNA Dataset

Protein sequences and their corresponding CDS sequences for three representative human viruses were downloaded from the NCBI database: Human coronavirus HKU1 (+ssRNA virus), Rotavirus A (dsRNA virus), and Human immunodeficiency virus (retroviral RNA). Due to the limited number of viral proteins and CDSs, all collected data were used as a test set to evaluate RNARL’s optimization capabilities for viral RNA types. As these viruses infect human hosts, the RNARL model trained on human mRNA data was employed for the optimization evaluations.

#### RNA Vaccine Patent Dataset

We retrieved two publicly available RNA vaccine patents (CN116019904A and CN117529335A) via Google Patents. Antigen protein sequences and their corresponding optimized mRNA sequences described in the patents were extracted for RNARL practical validation.

### RNARL Framework

The RNARL framework is an RNA codon design model based on generative models and reinforcement learning, designed to generate RNA sequences that translate into target proteins while exhibiting superior performance. The core of the RNARL framework is a reinforcement learning loop composed of an actor model (generator) and a reward model (evaluator) (Fig. 1).

The actor model generates candidate RNA sequences for a given protein sequence. Its architecture combines a Transformer Encoder with a MoE network. The protein sequences are first encoded into the embeddings using the pretrained protein language model ESM2_650M. These embeddings are then fed into the Transformer Encoder-MoE module, where they undergo multi-layer processing to generate nucleotide probability distributions at each position. Finally, the model generates RNA sequences via sampling.

The reward model evaluates the quality of generated RNA sequences. It takes RNA sequences as input, first encoding each nucleotide into a vector via a learned embedding layer. These vector sequences are then processed through a Transformer Encoder-MoE architecture similar to the actor model. Subsequently, a fully connected layer in the reward model generates quality scores, which serve as the reward signal during reinforcement learning training. Each RNA sequence is scored as follows:

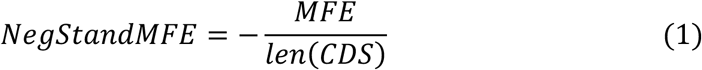

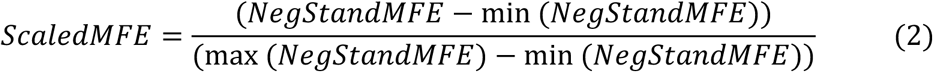

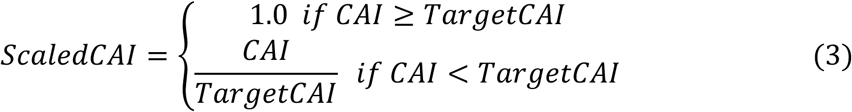

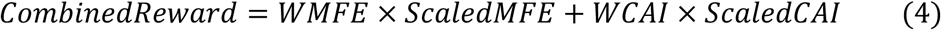

Here, *len*(*CDS*) denotes the length of CDS, *MFE* and *CAI* are the MFE and CAI values of CDS. *WMFE* , *WCAI* and *TargetCAI* are Hyperparameters. For the computational experiments presented in this study, we configured the following parameters: *WMFE* =2.0, *WCAI* =1.0, *TargetCAI* =0.8. The detailed selection process for these hyperparameters is presented in Supplementary Note 5 and Supplementary Fig. 6.

### GRPO Reinforcement Learning Algorithm

We employed the GRPO reinforcement learning algorithm to train the actor model to generate RNA sequences with high reward values based on input protein sequences. The GRPO algorithm achieves optimization of the actor network through policy gradient methods. GRPO incorporates the core principles of Proximal Policy Optimization (PPO), aiming to maximize rewards while maintaining policy stability by constraining the step size of policy updates. GRPO also introduces a KL divergence regularization term against a fixed reference policy (*π*_*θ*r*ef*_), as well as a mechanism for estimating the advantage function based on grouped sampling. Supplementary Note 4 and Supplementary Fig. 5 present the impact of GRPO reinforcement learning on the actor model.

#### GRPO Algorithm Objective Function

The core of the GRPO algorithm lies in maximizing an objective function that combines reward signals, policy stability constraints, and reference policy regularization. For a given state *s* and action *a*, the optimization objective is defined as:

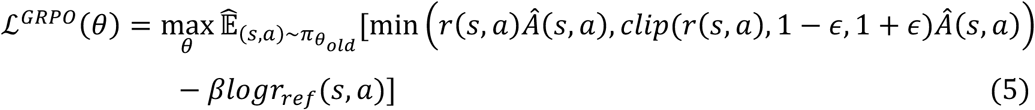

where 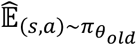 denotes the empirical expectation computed over data sampled under the old policy 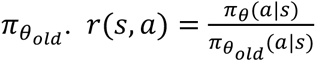 is the policy ratio between the current policy *π*_*θ*_ and the old policy 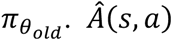 is the estimated advantage function for action *a* in state *s*. *clip*(*r*(*s*, *a*), 1 − *∈*, 1 + *∈*) truncates the policy ratio within the range [1 − *∈*, 1 + *∈*], where *∈* is the clipping coefficient (set to 0.2 in experiments). This term, borrowed from PPO’s design, restricts the magnitude of policy updates to prevent drastic policy changes. *logr*_*ref*_(*s*, *a*) = *logπ*_*θ*_(*a*|*s*) − *logπ*_*ref*_(*a*|*s*) represents the log policy ratio between the current policy *π*_*θ*_ and a fixed reference policy *π*_*ref*_ . This term is equivalent to a sample-based approximation of the KL divergence *D*_*KL*_(*π*_*θ*_||*π*_*ref*_). *β* (set to 0.01 in experiments) is the coefficient for the KL divergence regularization term, encouraging the current policy to remain close to the reference policy. The detailed selection process for these hyperparameters (*ε*, *β*) is presented in Supplementary Note 6 and Supplementary Fig. 7.

#### GRPO Training Procedure

In each training iteration, the GRPO algorithm executes the following steps to maximize the aforementioned objective function:

### Data sampling

For a batch of *N* input protein sequences 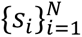, sample *G* candidate RNA sequences 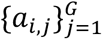 for each protein sequence *s*_*i*_ using the current old policy *π*_*θold*_ , where *G* is the group size. This results in a dataset of *N* × *G* (protein, RNA) pairs {(*s*_*i*_, *a*_*i*,j_)}_*i*=1…*N*,j=1…*G*_.

### Reward Evaluation

Compute the reward value *R*(*s*_*i*_, *a*_*i*,j_) for each sampled sequence (*s*_*i*_, *a*_*i*,j_).

### Advantage Function Estimation

For each protein sequence *s*_*i*_, compute the advantage function for its *G* sampled sequences by normalizing rewards within the group: Calculate the group mean 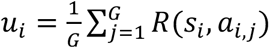 and standard deviation 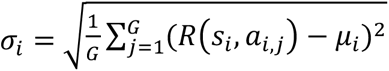 Compute the normalized advantage function 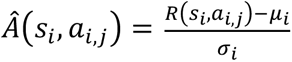.

### Objective Function Calculation and Policy Update

For each sampled pair (*s*_*i*_, *a*_*i*,j_), compute the probability ratios 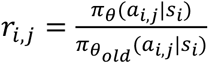 Compute the KL divergence approximation between the current policy and the fixed reference policy: *logr*_*ref*,*i*,j_ = *logπ*_*θ*_i*a*_*i*,j_l*s*_*i*_j − *logπ*_*θ*r*ef*_(*a*_*i*,j_|*s*_*i*_). Use the sampled data {(*s*_*i*_, *a*_*i*,j_)}, ratios *r*_*i*,j_, KL terms *logr*_*ref*,*i*,j_ , and advantages 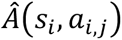 to compute the empirical objective function (a Monte Carlo approximation of the expectation in ℒ^*GRP*0^(*θ*)). Compute gradients of the objective function with respect to the actor model parameters *θ*. In practice, maximizing the objective is equivalent to minimizing its negative as a loss function. Update parameters *θ* via one step of gradient descent using an optimizer (AdamW).

### Old Policy Update

At fixed intervals (every 50 gradient steps in experiments), copy the current actor model parameters *θ* to the old policy parameters *θ*_*old*_ (*θ*_*old*_ ← *θ*) for subsequent sampling. The fixed reference policy parameters *θ*_*ref*_ remain unchanged throughout training.

### Length-Aware Distribution Sampling Technique

The training dataset exhibited a pronounced imbalance in protein sequence and RNA sequence length distribution. This imbalance inherently biased the model toward shorter sequences and compromised its performance on longer RNA sequences. To mitigate these issues, we implemented the LADS techniques. Protein sequences were binned into non-overlapping intervals of 100 amino acids (e.g., 1–100, 101–200, etc.), and adaptive sampling weights were assigned to each interval to equalize their contributions during training. Specifically, for each length bin, the sampling weight was calibrated inversely proportional to its occupancy, ensuring that the product of the weight and the number of proteins in each bin remained approximately constant across all intervals. This approach dynamically prioritizes underrepresented length categories during data sampling, thereby mitigating bias while preserving the diversity of sequences.

### Training Details

The actor and reward models were configured with the following hyperparameters: the Transformer Encoder utilized a hidden dimension of 768 and 8 layers, while the MoE component incorporated 6 experts with 2 experts activated per input token. Optimization was performed using the AdamW optimizer with a learning rate of 3.0 × 10^−B^, and the models were trained for 6 epochs to ensure convergence. Comprehensive hyperparameter tuning procedures are documented in the Supplementary Note 3 and Supplementary Table 1.

### Evaluation Metrics and Tools

We evaluated generated RNA sequences using the following metrics and bioinformatics tools:

#### MFE

Computed using LinearFold Package [60]. Lower MFE indicates higher structural stability. Normalized MFE (MFE / Sequence Length) is used in training to eliminate length bias.

#### CAI

Measures codon usage bias for species-specific translation efficiency. Codon usage bias information for each species was referenced from the Kazusa Codon Usage Database (https://www.kazusa.or.jp/codon/).

#### Proportion of G and C

Proportion of guanine (G) and cytosine (C) nucleotides.

#### Unpaired nt Proportion

Fraction of nucleotides in single-stranded regions.

Secondary structure prediction and analysis were performed using RNAfold [61]. This included the generation of the MFE structure, the positional entropy for each nucleotide position and the mountain plot representation of the MFE structure, the thermodynamic ensemble of RNA structures, and the centroid structure.

Tertiary structure prediction and analysis were performed using trRosettaRNA [62] to assess RNA compactness and stability.

### Baseline Implementation Details

The performance of the proposed RNARL was evaluated against five competing methods: GEMORNA [32], LinearDesign [15], CodonTransformer [5], a CAI-optimized method and a random method. For a fair and reproducible comparison, these five models were benchmarked by using their publicly available implementations and default configurations. The specific setup for each method is as follows:

GEMORNA: The evaluation was performed using the official implementation and pre-trained model weights from its GitHub repository (https://github.com/RainaBio/GEMORNA.git), with the provided inference scripts used without modification.

LinearDesign: The program was executed with its default parameter settings, using the software downloaded from its official repository (https://github.com/LinearDesignSoftware/LinearDesign.git).

CodonTransformer: Performance testing utilized the official code and pre-trained weights available on GitHub (https://github.com/Adibvafa/CodonTransformer.git).

CAI-optimized Method: This method involves the deterministic selection of the most frequent synonymous codon for each amino acid.

Random Method: This approach selects a synonymous codon for each amino acid through weighted random sampling, where weights are proportional to the natural codon usage frequencies of the target species.

### Web Interface

To enhance accessibility and facilitate the use of the RNARL framework, a user-friendly web interface was developed and hosted on Google Colab (https://colab.research.google.com/drive/1Z_t_Lt9CjqA0aygoNcdRzrThF1MOGBRb). This interface is designed to require no environment setup or programming knowledge, enabling users to readily input protein sequences and obtain optimized RNA sequences. The platform supports two modes of operation: the optimization of RNA sequences for a single protein by directly inputting its protein sequence, and the high-throughput optimization for multiple proteins simultaneously via the upload of CSV files. Additionally, users can select different species or RNA type model to perform targeted optimization based on their specific needs.

## Availability of Data and Materials

The multi-species mRNA and multi-type RNA datasets are available at https://zenodo.org/records/15486992. The source code is publicly accessible on GitHub (https://github.com/xlab-BioAI/RNARL) or Zenodo (https://doi.org/10.5281/zenodo.16931463) under the MIT license. An interactive web application developed and hosted on Google Colab is provided here (https://colab.research.google.com/drive/1Z_t_Lt9CjqA0aygoNcdRzrThF1MOGBRb), enabling users to efficiently optimize target RNA sequences.

## Acknowledgements

We thank the editor and the two anonymous reviewers for their inspiring and constructive comments and suggestions to greatly improve the quality and presentation of the work.

## Ethics declarations

### Ethics approval and consent to participate

Not applicable.

### Consent for publication

Not applicable.

### Authors’ Contributions

S.L. conceptualized the project, developed the model, performed the validation, and wrote the original draft. H.T., K.W., and R.W. participated in the literature review and model discussions. H.W. handled manuscript revision and acquired funding. T.Z. handled project administration and manuscript revision. Y.X. conceived and supervised the project, acquired funding, handled project administration, and revised the manuscript.

## Supplementary information

Supplementary information is available in a separate file accompanying this article.

