## supplementary information for "Reinforcement learning-driven unified generative framework for multi-objective RNA codon design"

**Table of Contents**


| **Supplementary Notes** | |
|---|---|
| Supplementary Note 1. Amino Acid Determinants of Mixture-of-Experts Activation | 3 |
| Supplementary Note 2. RNARL Enhances Structural Stability of RNA Vaccines | 3 |
| Supplementary Note 3. Hyperparameter Optimization of the Actor Model | 4 |
| Supplementary Note 4. Impact of Reinforcement Learning on the Actor Model | 4 |
| Supplementary Note 5. Hyperparameter Sensitivity Analysis of the RNARL Reward Function | 4 |
| Supplementary Note 6. GRPO Hyperparameter Sensitivity Analysis | 5 |
| Supplementary Note 7. Comparison of RNARL- and CodonTransformer-Generated Sequences in Similarity to Natural CDS | 6 |
| Supplementary Note 8. Impact of Clustering Parameters on Model Performance | 6 |
| Supplementary Note 9. Performance Comparison of a Unified Cross-Species Model and Species-Specific Models | 7 |
| Supplementary Note 10. Sequence-Level Determinants of RNARL Optimization Performance | 9 |
| **Supplementary Figures** | |
| Supplementary Fig. 1. Amino acid feature importance for expert activation in the MoE gating network. | 11 |
| Supplementary Fig. 2. Cross-species evaluation of RNA codon design methods. | 12 |
| Supplementary Fig. 3. Performance evaluation of codon design methods on retroviral RNA. | 12 |
| Supplementary Fig. 4. Comparative structural analysis of RNARL-optimized and patented RNA designs. | 13 |
| Supplementary Fig. 5. Performance improvement of the actor model during reinforcement learning. | 14 |
| Supplementary Fig. 6. Hyperparameter sensitivity analysis of the RNARL reward function. | 14 |
| Supplementary Fig. 7. Sensitivity analysis of GRPO hyperparameters. | 15 |
| Supplementary Fig. 8. Comparison of similarity to natural CDS for sequences generated by RNARL and CodonTransformer. | 17 |
| Supplementary Fig. 9. Impact of protein sequence clustering identity thresholds on model performance for the *Homo sapiens* dataset. | 17 |
| Supplementary Fig. 10. Performance comparison between the unified model and species-specific models. | 19 |
| Supplementary Fig. 11. The impact of protein properties on the optimization performance of the RNARL. | 19 |

| **Supplementary Tables** | |
|---|---|
| Supplementary Table 1. Hyperparameter optimization results for the actor model on the *Homo sapiens* dataset | 20 |
| Supplementary Table 2. The sizes of the training set, validation set, and test set for six different species mRNAs and human circular RNAs | 21 |
| Supplementary Table 3. Dataset sizes under different clustering identity thresholds | 21 |


### Supplementary Notes

#### Supplementary Note 1. Amino Acid Determinants of Mixture-of-Experts Activation

To investigate how different experts are activated within the Mixture-of-Experts (MoE), we examined the top 10 amino acid types that most significantly influence the activation of each expert, as depicted in Supplementary Fig. 1. According to Supplementary Fig. 1, different experts are predominantly activated by distinct amino acid types. For instance, Expert 2 is highly influenced by K (Lysine), Expert 4 by S (Serine), and Expert 5 by P (Proline). This suggests that different experts may specialize in processing sequences enriched in particular amino acid types. Furthermore, some experts are influenced by multiple amino acids. For example, both Expert 0 and Expert 1 show high importance for R (Arginine), P (Proline), and S (Serine), which may reflect the underlying biological relationships among these amino acids.

#### Supplementary Note 2. RNARL Enhances Structural Stability of RNA Vaccines

Supplementary Fig. 4 provides a comprehensive comparative analysis of the RNARL-optimized and patented RNA designs for encoding two distinct viral antigens, gL and gp42. Supplementary Fig. 4a and Supplementary Fig. 4b present predicted structural contact maps for the gL RNA sequence. For the RNARL-optimized sequence, these maps reveal an increased density of contact points compared to the patented design, suggesting the formation of more compact and potentially stable tertiary structures. Supplementary Fig. 4c and Supplementary Fig. 4d delve into comprehensive secondary structure characteristics for the gL RNA. These analyses, which include positional entropy and mountain plots illustrating the MFE, partition function ensemble, and centroid structures, provide insights into the thermodynamic landscape and typical folding patterns. This enhanced stability is further supported by the tertiary structure predictions for the gp42 RNA sequence shown in Supplementary Fig. 4e and Supplementary Fig. 4f, where the RNARL-optimized sequence folds into a demonstrably more compact and tightly packed conformation compared to the patent-provided sequence. In summary, these data elucidate how the RNARL optimization strategy influences various levels of RNA structure, from local contacts and secondary folding stability to global tertiary conformation.

**Supplementary Note 3. Hyperparameter Optimization of the Actor Model**

We performed hyperparameter optimization for the actor model on the *Homo sapiens* dataset, exploring six key parameters: the hidden dimension of Transformer Encoder (dmodel), the layers of Transformer Encoder (encoder_layers), the number of experts (num_experts), the number of experts activated per input token (top_k_experts), learning rate (learning_rate), and the number of epochs (num_epochs). The results of this hyperparameter search are presented in Supplementary Table 1. As observed in Supplementary Table 1, different hyperparameters had minimal impact on model performance, demonstrating the model's robustness. For our final configuration, we selected the following hyperparameter values: the Transformer Encoder utilized a hidden dimension of 768 and 8 layers, while the Mixture-of-Experts (MoE) component incorporated 6 experts with 2 experts activated per input token. Optimization was performed using the AdamW optimizer with a learning rate of $3.0\times10^{-5}$, and the model was trained for 6 epochs to ensure convergence.

**Supplementary Note 4. Impact of Reinforcement Learning on the Actor Model**

To quantify the impact of reinforcement learning, we evaluated the actor model's performance throughout the training epochs. The results (Supplementary Fig. 5) demonstrate improvements in key metrics after six epochs: the median CAI increased by 0.015, and the median MFE decreased from -321.7 kcal/mol to -337.35 kcal/mol. These gains affirm that the reinforcement learning component effectively guides the actor model toward generating sequences with enhanced properties, thereby validating the multi-objective optimization capability of the RNARL framework.

**Supplementary Note 5. Hyperparameter Sensitivity Analysis of the RNARL Reward Function**

To systematically select the reward function hyperparameters in RNARL, we performed a series of sensitivity analyses. First, for the TargetCAI hyperparameter, we evaluated four candidate values (0.6, 0.7, 0.8, and 0.9). Next, we examined the WMFE and WCAI hyperparameters, which control the relative contributions of MFE and CAI in the optimization objective. For this analysis, we fixed WCAI at 1.0 and varied WMFE to assess its influence on model performance. The results are summarized in Supplementary Fig. 6.

As shown in Supplementary Fig. 6a, increasing WMFE led to slight improvements in the MFE of the generated RNA sequences. However, when WMFE was increased to 3.0, a substantial decrease in CAI values was observed. WMFE = 2.0 was therefore chosen as a trade-off that improves MFE while avoiding a major reduction in translational efficiency. The results in Supplementary Fig. 6b indicate that the TargetCAI hyperparameter has a strong influence on the final CAI of the generated sequences. A clear improvement in CAI was observed when TargetCAI was increased from 0.7 to 0.8. This choice is further supported by the prevailing view in the field that a CAI of ≥0.8 is indicative of high translational efficiency. Accordingly, TargetCAI was set to 0.8 in the RNARL framework.

**Supplementary Note 6. GRPO Hyperparameter Sensitivity Analysis**

To assess the influence of GRPO hyperparameters on model performance, we performed a sensitivity analysis for the KL divergence coefficient $\beta$ and the GRPO clipping coefficient $\varepsilon$. The results are shown in Supplementary Fig. 7.

Supplementary Fig. 7a shows that while increasing the hyperparameter $\beta$ slightly reduces minimum CAI values, its overall effect on model performance is minimal. We selected $\beta$ = 0.01 as it provides a favorable trade-off between CAI and MFE. Although its minimum CAI is slightly lower than that at $\beta$ = 0.005, it yields better MFE values for longer sequences than the other $\beta$ settings. The clipping parameter $\varepsilon$ has a more substantial impact on the CAI values (Supplementary Fig. 7b). The model generates sequences with significantly better CAI scores when $\varepsilon$ is set to 0.15 or 0.2, compared to when $\varepsilon$ is 0.1 or 0.25. We ultimately chose $\varepsilon$ = 0.2. This value was selected because it yielded the best CAI performance while maintaining excellent MFE performance.

**Supplementary Note 7. Comparison of RNARL- and CodonTransformer-Generated Sequences in Similarity to Natural CDS**

We compared RNA sequences generated by RNARL and CodonTransformer with their corresponding natural sequences in terms of sequence similarity. Two metrics were used: nucleotide sequence identity and codon-usage similarity. Nucleotide sequence identity measures the fraction of identical nucleotides at aligned positions between two sequences. Codon-usage similarity was computed by first deriving the codon-frequency distribution for each sequence and then quantifying the similarity between the two distributions as 1-JSD (Jensen–Shannon divergence). Because JSD ranges from 0 to 1, with smaller values indicating more similar distributions, 1-JSD also ranges from 0 to 1, with larger values indicating higher codon-usage similarity.

Across both metrics, a consistent pattern was observed. For all species, sequences generated by both RNARL and CodonTransformer exhibit >70% similarity to the corresponding natural CDS (Supplementary Fig. 8). Except for *Thermococcus kodakarensis KOD1*, CodonTransformer generally produces sequences that are more similar to natural CDS than those generated by RNARL. This is expected, as CodonTransformer is pre-trained on natural sequences with the explicit objective of modeling their distribution, whereas RNARL is optimized to improve sequence properties such as higher CAI and lower MFE, rather than to closely reproduce natural coding sequences.

**Supplementary Note 8. Impact of Clustering Parameters on Model Performance**

To evaluate how clustering parameters affect model performance, we conducted computational experiments on the *Homo sapiens* dataset. Protein sequences were clustered under three identity thresholds: (1) 30% identity over 80% length, (2) 60% identity over 80% length, and (3) 90% identity over 80% length. For each condition, one representative sequence per cluster was selected, and the resulting dataset was split into training (90%), validation (5%), and test (5%) sets. The dataset sizes are summarized in Supplementary Table 3.

We trained and evaluated RNARL separately on each clustered dataset and re-evaluated all benchmark methods on the corresponding test sets (Supplementary Fig. 9). For RNARL, the identity threshold mainly affected CAI (Supplementary Fig. 9a). The mean CAI increased from 0.927 ± 0.047 at 30% identity to 0.943 ± 0.044 at 60% identity and 0.967 ± 0.027 at 90% identity. Notably, even at 30% and 60% identity, RNARL-generated sequences achieved higher CAI values than all state-of-the-art methods except the CAI-optimized baseline (Supplementary Fig. 9b, c), indicating strong generalization.

The effect on MFE was less pronounced but followed a similar trend (Supplementary Fig. 9a). The mean MFE was −581.4 ± 370.1 kcal/mol at 30% identity, −607.5 ± 379.1 kcal/mol at 60% identity, and −623.4 ± 396.3 kcal/mol at 90% identity. Even at 30% and 60% identity, RNARL's MFE performance remained superior to all methods except LinearDesign. Overall, as clustering becomes more stringent, the performance of RNARL decreases slightly. Despite this, RNARL remains highly competitive and generally outperforms state-of-the-art methods across all clustering settings, underscoring the robustness of the framework.

### Supplementary Note 9. Performance Comparison of a Unified Cross-Species Model and Species-Specific Models

To compare a unified modeling strategy with species-specific approaches, we constructed a cross-species unified model and evaluated its performance against six individually trained, species-specific models. For the unified model, we jointly trained it on datasets from all six species with minimal modifications to the original architecture. The model was conditioned on a 64-dimensional vector of species-specific codon frequencies, which influences both sequence generation and scoring. For the actor model, this vector is passed through a three-layer fully connected network (FCN) to produce a weight vector, which is then log-transformed and added to the output logits. For the reward model, the same FCN architecture maps the frequency vector to a feature vector, which is fused with the encoder outputs at each token position via a residual connection.

We then compared RNA sequences generated by the unified model and the six specialized models across multiple metrics (Supplementary Fig. 10). Overall, the two approaches exhibited comparable performance, with no consistent advantage for either.

For CAI, the relative performance of the two strategies varied by species. The species-specific models achieved higher CAI values for *Homo sapiens*, *Thermococcus kodakarensis KOD1*, and *Arabidopsis thaliana*, whereas the unified model yielded higher CAI values for *Chlamydomonas reinhardtii* and *Penicillium chrysogenum*. The specialized models for *Homo sapiens* and *Arabidopsis thaliana* likely benefit from larger training datasets, enabling these models to capture species-specific sequence patterns more effectively. Similarly, *Thermococcus kodakarensis KOD1* possesses a highly distinct codon usage bias adapted to extreme temperatures. This pattern can be learned more precisely by a specialized model without interference from other species. Conversely, for species with smaller datasets (*Chlamydomonas reinhardtii*, *Penicillium chrysogenum*), the unified model likely outperforms the specialized models by leveraging cross-species transfer learning from the larger, aggregated dataset.

For MFE, the differences between the two approaches were minor. The species-specific models achieved slightly lower (more favorable) MFE values for *Chlamydomonas reinhardtii* and *Penicillium chrysogenum*, whereas the unified model yielded lower MFE values for Arabidopsis thaliana, *Thermococcus kodakarensis KOD1*, and *Escherichia coli*. The limited variation in MFE is expected, as the biophysical principles governing RNA folding are universal and not species-dependent. Both model types effectively learn to generate structurally stable sequences. The minor performance differences observed are likely secondary effects of the different codon choices made to optimize species-specific CAI.

In conclusion, these results indicate that, when conditioned on codon usage frequencies, the unified model can effectively learn and apply distinct codon preferences across multiple species. Its performance is comparable to that of individually trained, species-specific models. This finding supports the interpretation that CAI optimization is the primary learning signal, suggesting that model performance is driven more by codon usage bias than by differences in species-specific amino acid distributions.

**Supplementary Note 10. Sequence-Level Determinants of RNARL Optimization**

**Performance**

To analyze sequence-level optimization performance, we defined two core 'optimization gain' metrics. The first, normalized MFE gain, is computed as the difference between the MFE of the natural and optimized RNA, divided by RNA length. Because lower MFE corresponds to more stable RNA secondary structures, larger positive values reflect greater stability gains, and length normalization reduces bias arising from sequence length. The second metric, CAI gain, is defined as the CAI of the optimized RNA minus that of the natural RNA. Standard bioinformatics tools (Biopython) were used to calculate physicochemical properties for each protein on the *Homo sapiens* dataset, including hydrophobicity, positive charge, aromaticity, isoelectric point, instability index and helix propensity.

A global analysis of the relationships between optimization gains and protein features was first performed using a Spearman correlation heatmap (Supplementary Fig. 11a). In particular, normalized MFE gain and CAI gain show a strong positive correlation ($r = 0.64$, $p < 0.01$), indicating that RNARL can simultaneously improve RNA stability and codon usage efficiency. To further illustrate this relationship, an optimization trade-off landscape was constructed (Supplementary Fig. 11b) using CAI gain and normalized MFE gain. Most points cluster in the upper-right quadrant, where both gains are positive, indicating that RNARL effectively achieves multi-objective optimization.

Further investigation focused on specific negative associations identified in the heatmap. For instance, the relationship between CAI gain and the protein instability index (Supplementary Fig. 11c) shows a clear downward trend ($r = -0.21$, $p < 0.01$), indicating that intrinsically more unstable proteins tend to exhibit smaller improvements in codon adaptation. Analysis of the relationship between CAI gain and protein hydrophobicity (Supplementary Fig. 11d) revealed a weak positive correlation ($r = 0.19$, $p < 0.01$), suggesting that more hydrophobic proteins achieve slightly larger CAI gains, although hydrophobicity alone is not a major determinant of optimization gains. To explore potential non-linear effects, a stratified analysis was performed on the normalized MFE gain based on protein hydrophobicity (Supplementary Fig. 11e). In this analysis, proteins were divided into quartiles according to their hydrophobicity, and the distribution of normalized MFE gain was examined within each group. The stratified analysis reveals a subtle pattern: as hydrophobicity increases, the median normalized MFE gain shows a slight downward trend. Finally, an integrated view of ‘optimizability’ was established by defining a composite optimization score as the sum of normalized MFE gain and CAI gain. Based on this score, proteins were categorized into ‘easy-to-optimize’ (top 10%) and ‘hard-to-optimize’ (bottom 10%) groups. The ‘hard-to-optimize’ proteins display lower hydrophobicity but higher instability than their ‘easy-to-optimize’ counterparts (Supplementary Fig. 11f).

In summary, the sequence-level analysis demonstrates that RNARL effectively achieves concomitant improvements in RNA stability and codon adaptation. More importantly, the magnitude of these optimization gains is not uniform but is instead governed by the intrinsic physicochemical properties of the encoded protein. By integrating these gain metrics into a composite score, we successfully delineate a ‘hard-to-optimize’ protein subset, characterized by a distinct signature of low hydrophobicity and high instability. These findings offer critical insights into the biophysical constraints of sequence optimization.

### Supplementary Figures

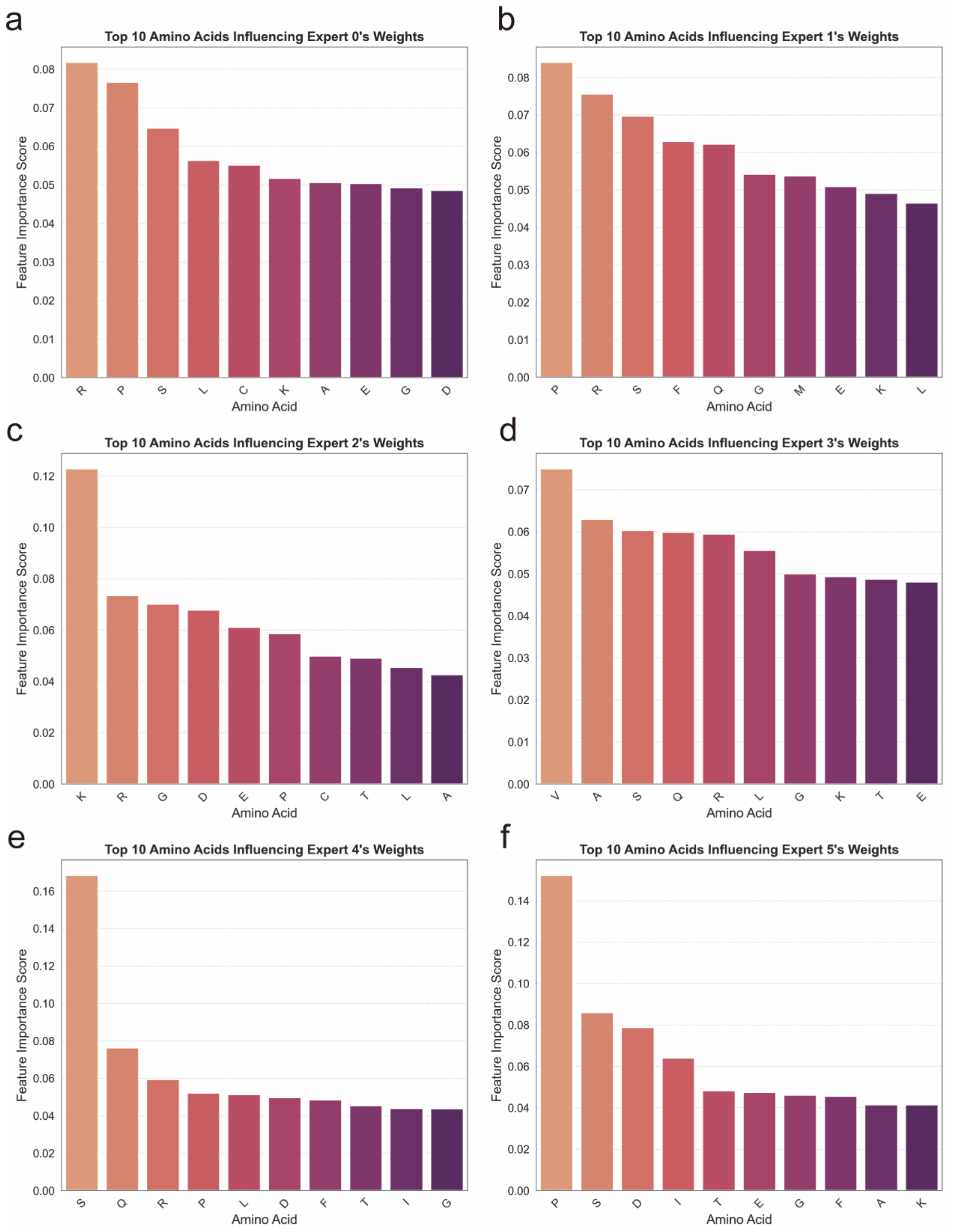


**Supplementary Fig. 1.** Amino acid feature importance for expert activation in the MoE gating network. **a-f**, the top 10 amino acids influencing the activation status (weights) of Expert 0 (a), Expert 1 (b), Expert 2 (c), Expert 3 (d), Expert 4 (e), and Expert 5 (f) within the MoE gating network. Feature importance scores quantify the influence of each amino acid on the respective expert's activation.

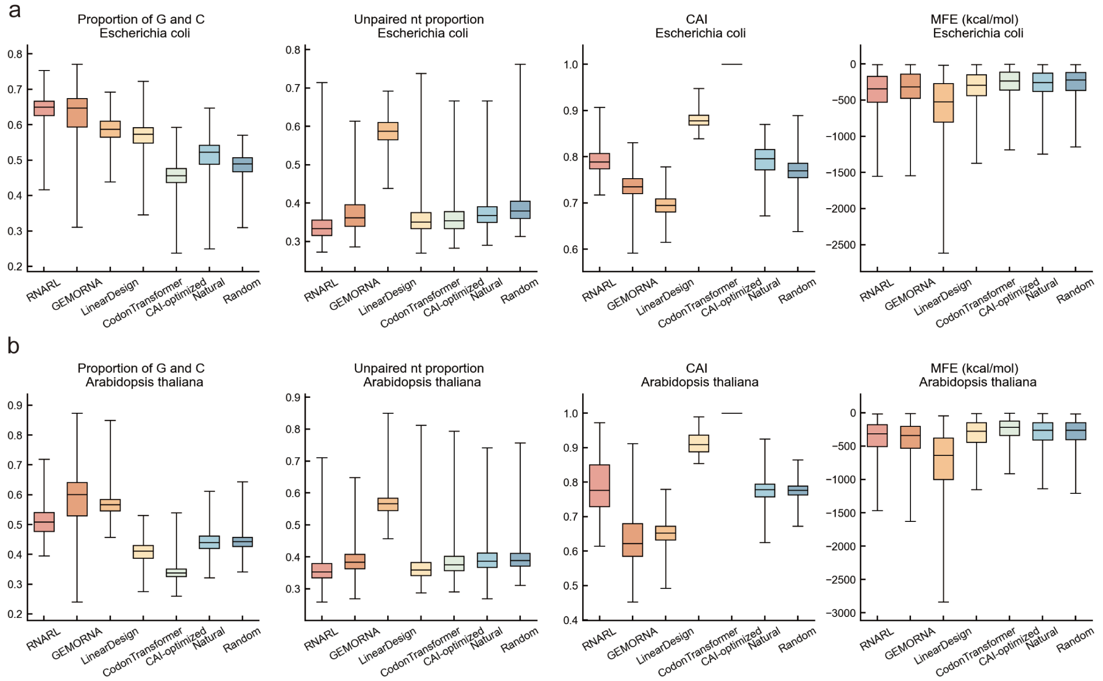


**Supplementary Fig. 2.** Cross-species evaluation of RNA codon design methods. **a,** Results for *Escherichia coli*. **b,** Results for *Arabidopsis thaliana*.

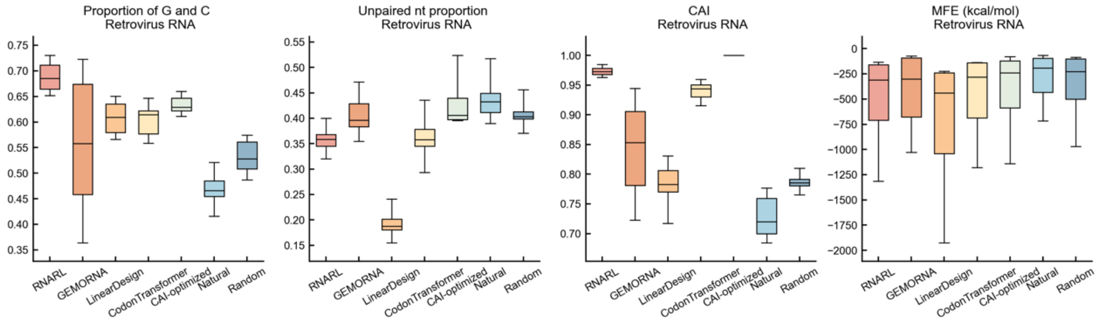


**Supplementary Fig. 3.** Performance evaluation of codon design methods on retroviral RNA.

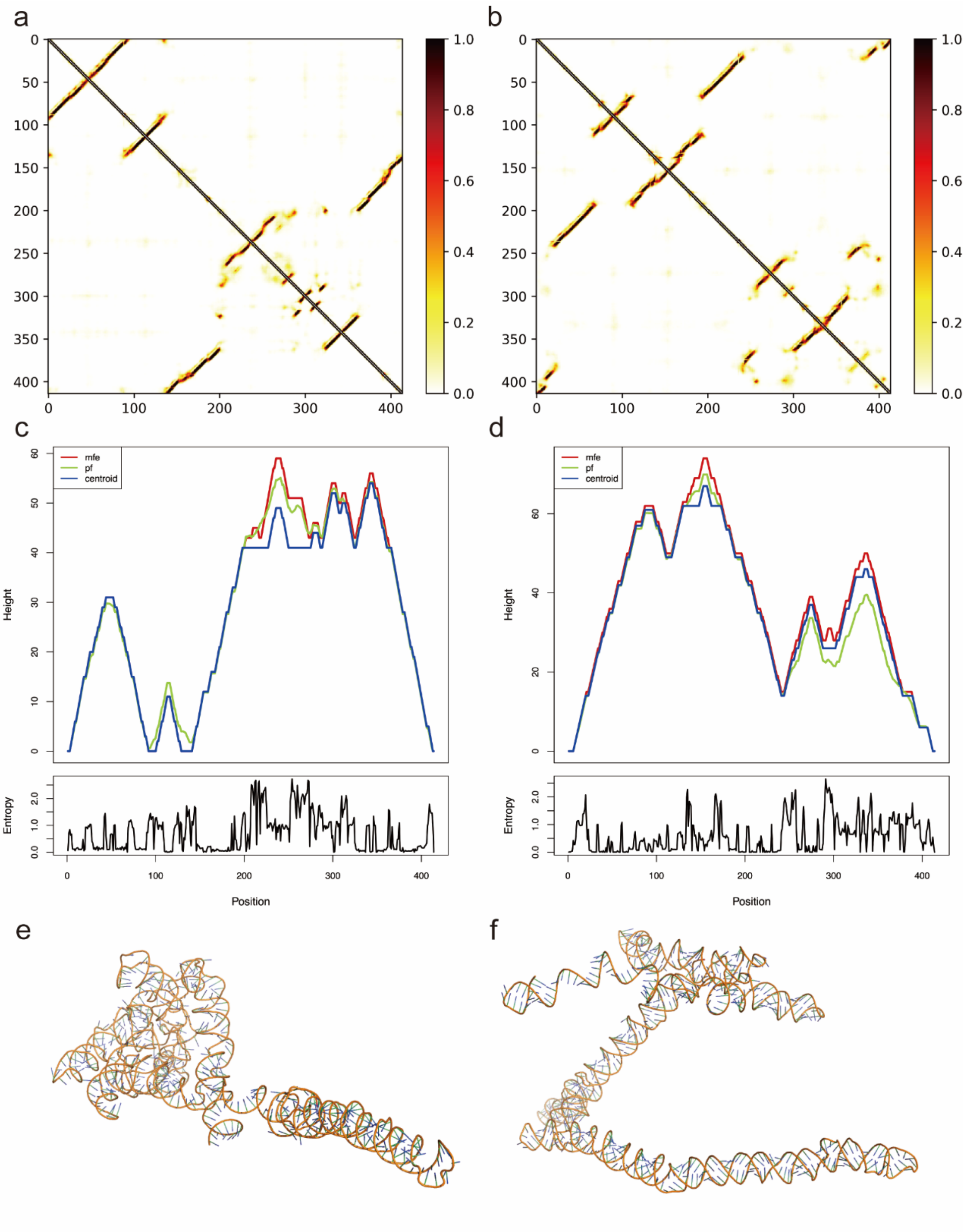


**Supplementary Fig. 4.** Comparative structural analysis of RNARL-optimized and patented RNA designs. **a, b,** Structural contact maps for the RNA sequence encoding gL, showing the RNARL-optimized design (a) and the patented design (b). The heatmap scale indicates contact probability from 0 (white) to 1 (dark red). **c, d,** Comprehensive secondary structure analysis for the gL RNA sequences. Mountain plots for the RNARL-optimized design (c) and the patented design (d) illustrate the MFE structure (red line), the partition function (pf) ensemble (green line), and the centroid structure (blue line). The lower panels display positional entropy across the sequence.

**e, f,** Tertiary structures of the RNA sequence encoding gp42, comparing the RNARL-optimized design (e) with the patented design (f).

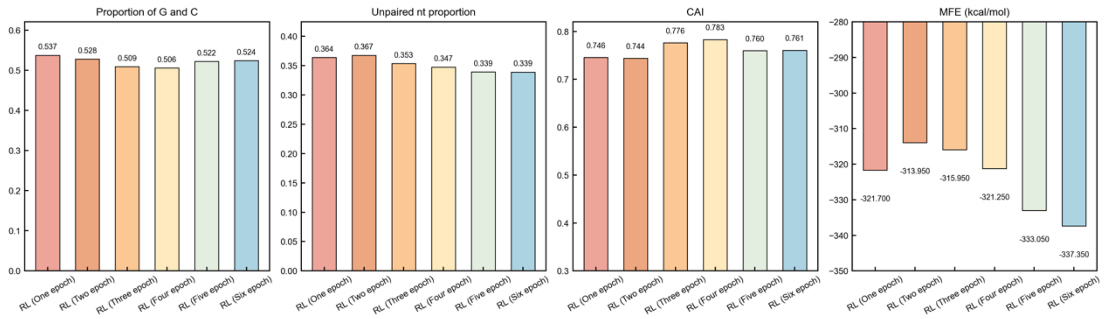


**Supplementary Fig. 5.** Performance improvement of the actor model during reinforcement learning. The plots show the median CAI and MFE evaluated on the *Arabidopsis thaliana* test set after each training epoch.

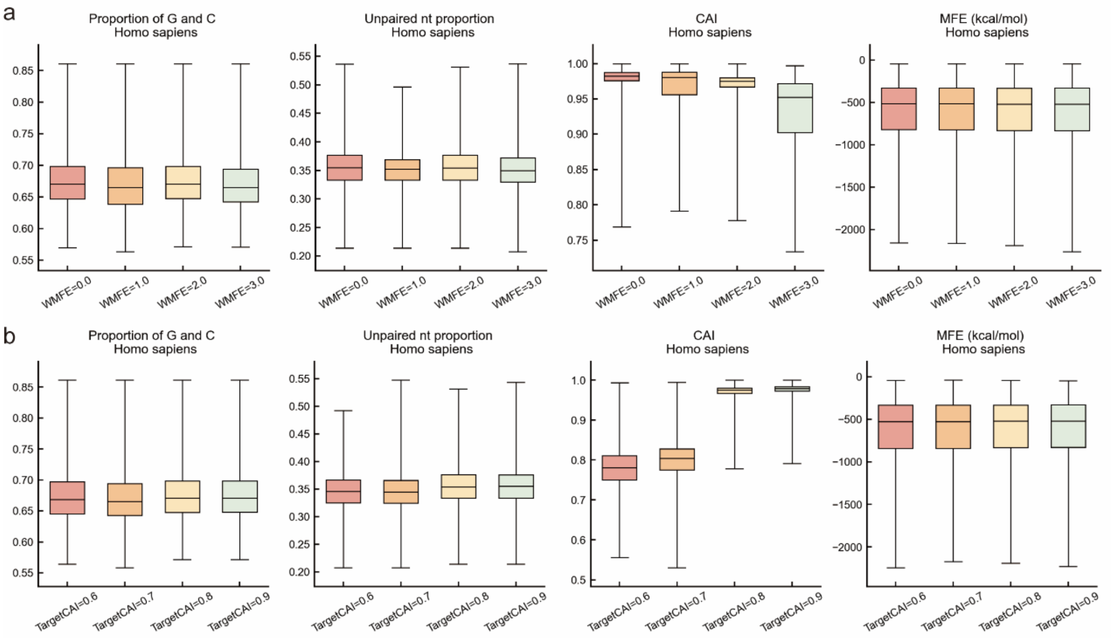


**Supplementary Fig. 6.** Hyperparameter sensitivity analysis of the RNARL reward function. Box plots show the distributions of four key RNA properties of sequences. (a) Effect of varying WMFE from 0.0 to 3.0 while holding WCAI constant. (b) Effect of varying TargetCAI from 0.6 to 0.9.

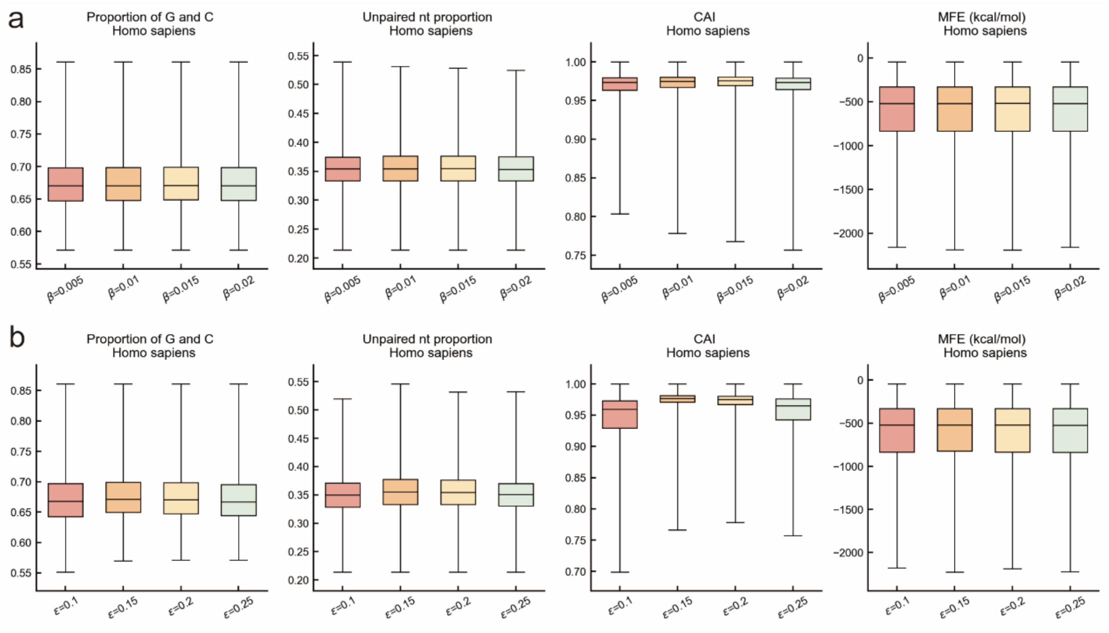


**Supplementary Fig. 7**. Sensitivity analysis of GRPO hyperparameters. Box plots showing the distributions of four key properties of RNA sequences. **a**. Effect of varying the KL divergence coefficient $\beta$. **b**. Effect of varying the GRPO clipping coefficient $\varepsilon$.

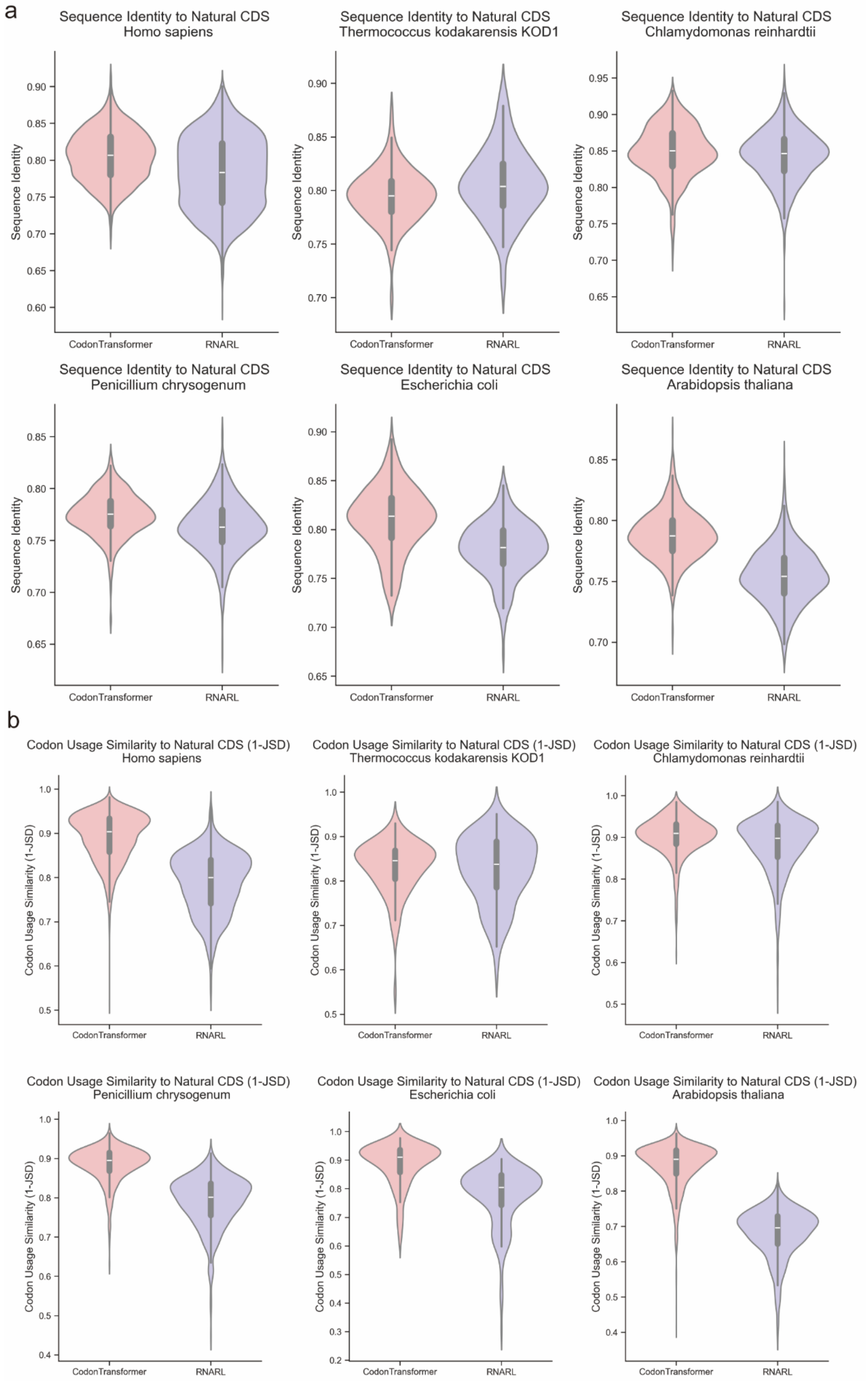
a
Sequence Identity to Natural CDS
Homo sapiens
Sequence Identity to Natural CDS
Thermococcus kodakarensis KOD1
Sequence Identity to Natural CDS
Chlamydomonas reinhardtii
Sequence Identity to Natural CDS
Penicillium chrysogenum
Sequence Identity to Natural CDS
Escherichia coli
Sequence Identity to Natural CDS
Arabidopsis thaliana
Sequence Identity
CodonTransformer
RNARL
b
Codon Usage Similarity to Natural CDS (1-JSD)
Homo sapiens
Codon Usage Similarity to Natural CDS (1-JSD)
Thermococcus kodakarensis KOD1
Codon Usage Similarity to Natural CDS (1-JSD)
Chlamydomonas reinhardtii
Codon Usage Similarity to Natural CDS (1-JSD)
Penicillium chrysogenum
Codon Usage Similarity to Natural CDS (1-JSD)
Escherichia coli
Codon Usage Similarity to Natural CDS (1-JSD)
Arabidopsis thaliana
Codon Usage Similarity (1-JSD)
CodonTransformer
RNARL

**Supplementary Fig. 8.** Comparison of similarity to natural CDS for sequences generated by RNARL and CodonTransformer.

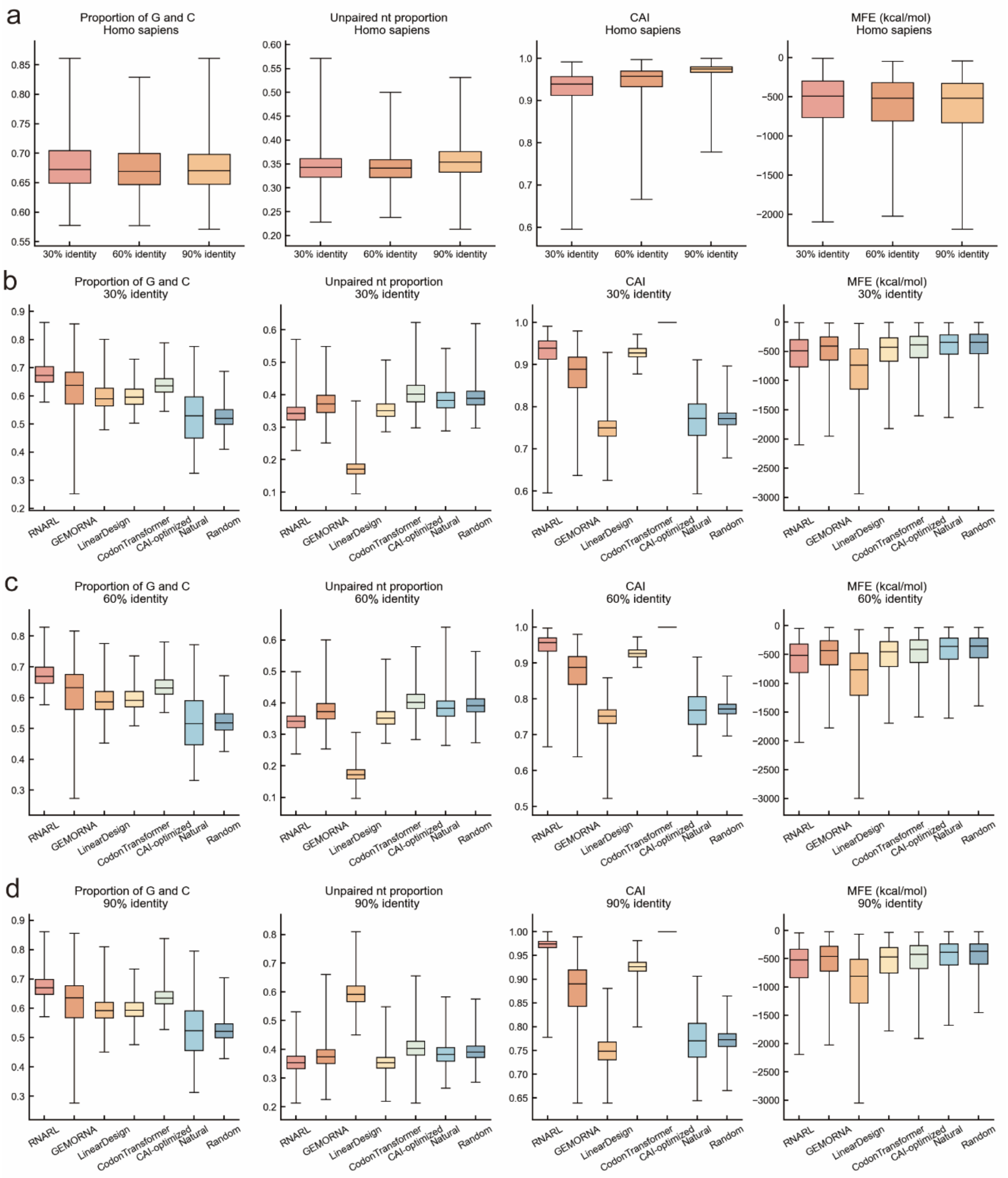


**Supplementary Fig. 9.** Impact of protein sequence clustering identity thresholds on model performance for the *Homo sapiens* dataset. **a,** Performance of RNARL trained on datasets derived from 30%, 60%, and 90% identity clustering. Box plots show the distribution of four key metrics for sequences generated by RNARL. **b-d,** Comparative performance of all codon design methods on test sets derived from 30% identity (**b**), 60% identity (**c**) and 90% identity (**d**) clustering, respectively.

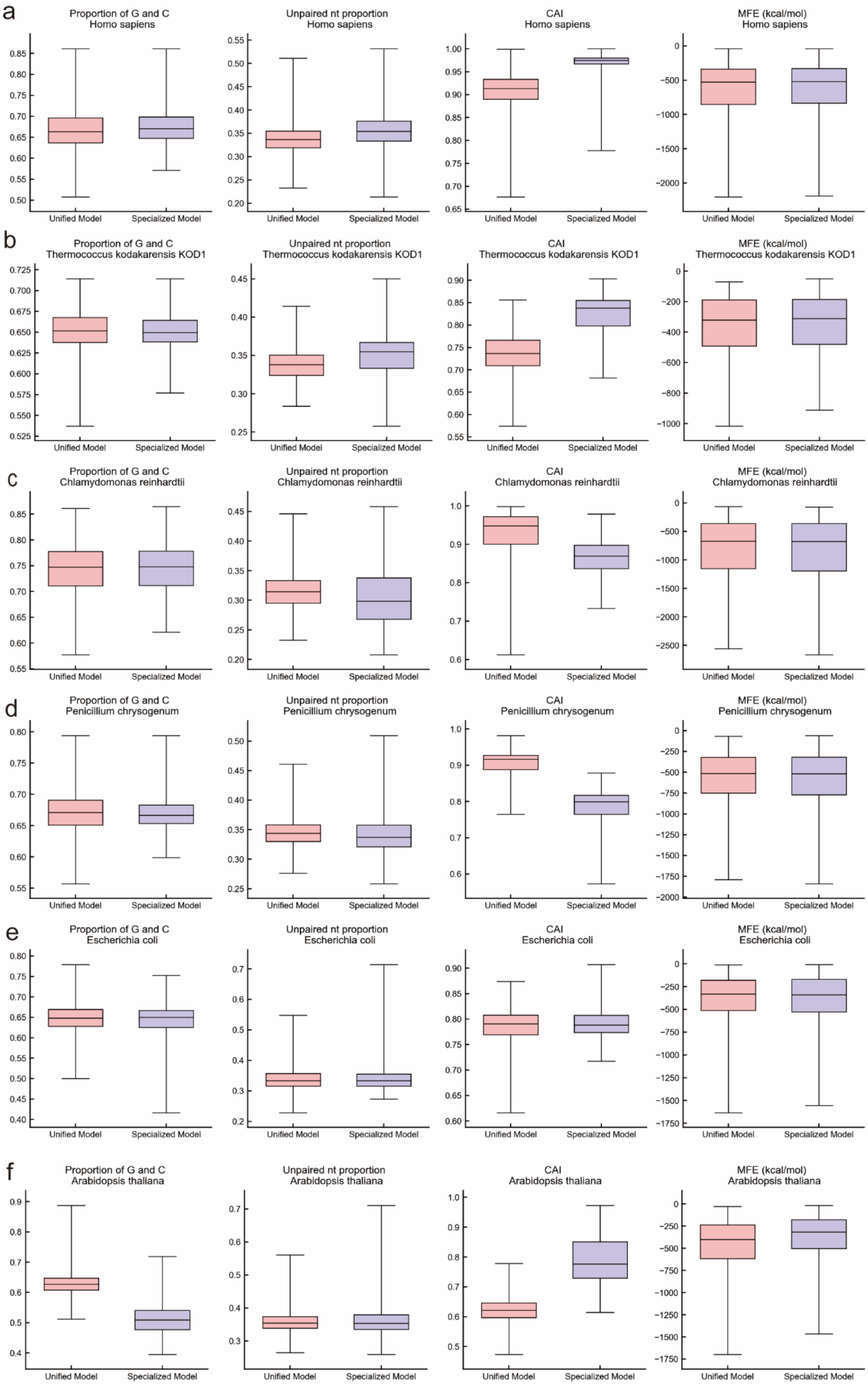
a
Proportion of G and C
Homo sapiens
Unpaired nt proportion
Homo sapiens
CAI
Homo sapiens
MFE (kcal/mol)
Homo sapiens
b
Proportion of G and C
Thermococcus kodakarensis KOD1
Unpaired nt proportion
Thermococcus kodakarensis KOD1
CAI
Thermococcus kodakarensis KOD1
MFE (kcal/mol)
Thermococcus kodakarensis KOD1
c
Proportion of G and C
Chlamydomonas reinhardtii
Unpaired nt proportion
Chlamydomonas reinhardtii
CAI
Chlamydomonas reinhardtii
MFE (kcal/mol)
Chlamydomonas reinhardtii
d
Proportion of G and C
Penicillium chrysogenum
Unpaired nt proportion
Penicillium chrysogenum
CAI
Penicillium chrysogenum
MFE (kcal/mol)
Penicillium chrysogenum
e
Proportion of G and C
Escherichia coli
Unpaired nt proportion
Escherichia coli
CAI
Escherichia coli
MFE (kcal/mol)
Escherichia coli
f
Proportion of G and C
Arabidopsis thaliana
Unpaired nt proportion
Arabidopsis thaliana
CAI
Arabidopsis thaliana
MFE (kcal/mol)
Arabidopsis thaliana
Unified Model
Specialized Model

**Supplementary Fig. 10.** Performance comparison between a unified model and six species-specific models. Box plots show the distribution of key metrics for RNA sequences generated by the unified model (trained on all six species) and the specialized models (trained individually for each species). **a-f** correspond to the six species: (**a**) *Homo sapiens*, (**b**) *Thermococcus kodakarensis KOD1*, (**c**) *Chlamydomonas reinhardtii*, (**d**) *Penicillium chrysogenum*, (**e**) *Escherichia coli*, and (**f**) *Arabidopsis thaliana*.

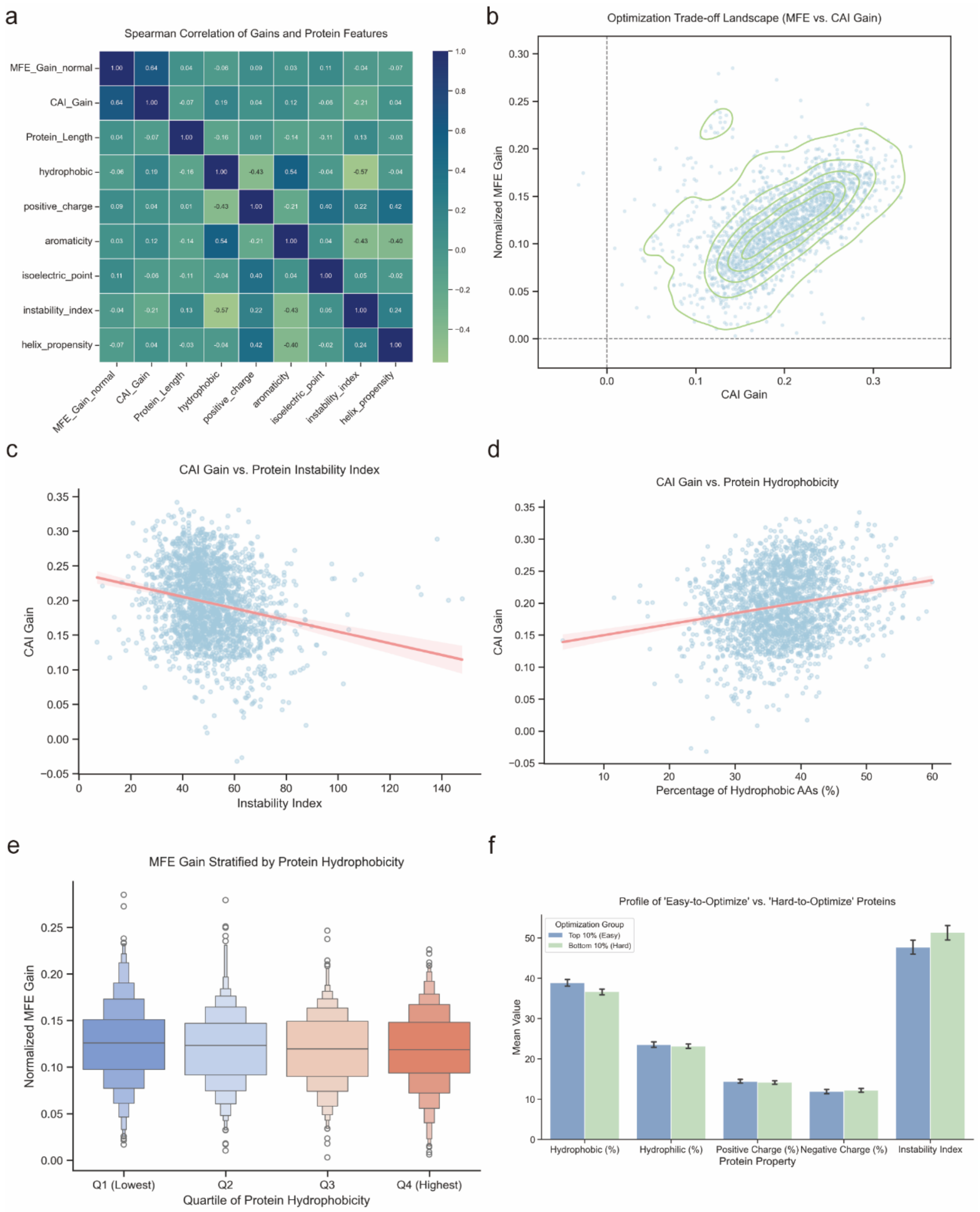


**Supplementary Fig. 11.** The impact of protein properties on the optimization performance of the RNARL model. **a,** Spearman correlation between optimization gains and protein physicochemical properties. **b,** Joint landscape of normalized MFE gain and CAI gain. **c,** Negative correlation between CAI gain and protein Instability index. **d,** Positive correlation between CAI gain and the fraction of hydrophobic amino acids. **e,** Influence of protein hydrophobicity on normalized MFE gain. **f,** Comparison of physicochemical properties between ‘easy-to-optimize’ and ‘hard-to-optimize’ proteins.

### Supplementary Tables

**Supplementary Table 1. Hyperparameter optimization results for the actor model on the *Homo sapiens* dataset**

| Hyperparameter | Value | Train loss | Val loss |
|---|---|---|---|
| dmodel | 256 | 2.204957 | 2.092456 |
| dmodel | 512 | 2.101884 | 2.088353 |
| dmodel | 640 | 2.051176 | 2.087566 |
| dmodel | 768 | 2.012526 | 2.086795 |
| encoder_layers | 5 | 2.003237 | 2.086227 |
| encoder_layers | 6 | 2.005026 | 2.086155 |
| encoder_layers | 7 | 2.008675 | 2.086072 |
| encoder_layers | 8 | 2.012526 | 2.086795 |
| num_experts | 3 | 2.015444 | 2.086109 |
| num_experts | 4 | 2.013328 | 2.087157 |
| num_experts | 5 | 2.011705 | 2.085867 |
| num_experts | 6 | 2.012526 | 2.086795 |
| top_k_experts | 1 | 2.020367 | 2.087599 |
| top_k_experts | 2 | 2.012526 | 2.086795 |
| top_k_experts | 3 | 2.016369 | 2.087683 |
| top_k_experts | 4 | 2.025556 | 2.087520 |
| learning_rate | 1.5e-5 | 2.138721 | 2.089686 |
| learning_rate | 2.0e-5 | 2.086384 | 2.088249 |
| learning_rate | 2.5e-5 | 2.040630 | 2.088372 |
| learning_rate | 3.0e-5 | 2.012526 | 2.086795 |
| num_epochs | 3 | 2.012526 | 2.086795 |

| | | | |
|---|---|---|---|
| num_epochs | 4 | 1.979588 | 2.085535 |
| num_epochs | 5 | 1.967966 | 2.084957 |
| num_epochs | 6 | 1.907094 | 2.083137 |

**Supplementary Table 2. The sizes of the training set, validation set, and test set for six different species mRNAs and human circular RNAs**

| Dataset type | Homo sapiens | Arabidopsis thaliana | Escherichia coli | Penicillium chrysogenum |
|---|---|---|---|---|
| Train Dataset | 38139 | 28144 | 3764 | 10382 |
| Validation Dataset | 2119 | 1564 | 209 | 577 |
| Test Dataset | 2119 | 1564 | 210 | 577 |
| | Thermococcus kodakarensis KOD1 | Chlamydomonas reinhardtii | Human circRNA | |
| Train Dataset | 1720 | 13653 | 21533 | |
| Validation Dataset | 96 | 759 | 1197 | |
| Test Dataset | 96 | 759 | 1196 | |

**Supplementary Table 3**. ***Homo sapiens*** **Dataset sizes under different clustering identity thresholds**

| Identity threshold | Train Dataset | Validation Dataset | Test Dataset |
|---|---|---|---|
| 30% | 24909 | 1384 | 1384 |
| 60% | 26156 | 1454 | 1453 |
| 90% | 38139 | 2119 | 2119 |
